# Elevated CD47 is a hallmark of dysfunctional aged muscle stem cells that can be targeted to augment regeneration

**DOI:** 10.1101/2022.04.29.489435

**Authors:** Ermelinda Porpiglia, Thach Mai, Peggy Kraft, Colin A. Holbrook, Antoine de Morree, Veronica D. Gonzalez, Keren Hilgendorf, Laure Fresard, Angelica Trejo, Sriram Bhimaraju, Peter K. Jackson, Wendy J. Fantl, Helen M. Blau

## Abstract

In aging, skeletal muscle strength and regenerative capacity declines due, in part, to functional impairment of muscle stem cells (MuSCs), yet the underlying mechanisms remain elusive. Here we capitalize on mass-cytometry to identify high CD47 expression as a hallmark of dysfunctional MuSCs (CD47^hi^) with impaired regenerative capacity that predominate with aging. The prevalent CD47^hi^ MuSC subset suppresses the residual functional CD47^lo^ MuSC subset through a paracrine signaling loop, leading to impaired proliferation. We uncover that elevated CD47 levels on aged MuSCs result from increased U1 snRNA expression, which disrupts alternative polyadenylation. The deficit in aged MuSC function in regeneration can be overcome either by morpholino-mediated blocking of CD47 alternative polyadenylation or antibody blockade of CD47 signaling, leading to improved regeneration in aged mice, with therapeutic implications. Our findings highlight a previously unrecognized age-dependent alteration in CD47 levels and function in MuSCs, which underlies reduced muscle repair in aging.

## INTRODUCTION

Skeletal muscles make up 40% of the body’s mass. After the age of 50, humans lose on average 15% of their muscle mass per decade, culminating in the drastic loss of muscle strength characteristic of sarcopenia(von Haehling et al., 2010). The loss of strength with aging leads to diminished autonomy in the elderly and is associated with risk factors for disabling conditions, such as osteoporosis, heart failure, and cognitive decline(Martinez et al., 2015; Robertson et al., 2013; Rockwood and Mitnitski, 2007). There are currently no therapies for sarcopenia and its financial burden is high, at approximately $19 billion in annual healthcare costs in the United States alone(Goates et al., 2019; Rolland et al., 2008). Sarcopenia is associated with the loss of functional skeletal muscle stem cells (MuSCs). MuSCs, also known as satellite cells, reside within skeletal muscle tissue in niches juxtaposed to myofibers and are required for skeletal muscle maintenance and regeneration throughout life(Blau et al., 2015; Feige et al., 2018; Keefe et al., 2015; MAURO, 1961; de Morree et al., 2019; Pawlikowski et al., 2015; Schmidt et al., 2019; Sousa-Victor and Muñoz-Cánoves, 2016). Changes in cell extrinsic regulators such as fibronectin, wnt, fibroblast growth factor-2 (FGF-2), and apelin in the muscle microenvironment diminish MuSC function with aging(Chakkalakal et al., 2012; Lukjanenko et al., 2016, 2019; Rozo et al., 2016; Vinel et al., 2018). In addition MuSCs isolated from aged mice exhibit intrinsic defects due to aberrant p38 MAPK, JAK/STAT and TGF-*β*signaling, which lead to a decrease in the proportion of functional MuSCs, hindering muscle regeneration(Bernet et al., 2014; Brett et al., 2020; Cosgrove et al., 2014; Price et al., 2014; Tierney et al., 2014). The absence of markers for distinguishing and prospectively isolating dysfunctional MuSCs has limited mechanistic insights and the development of therapeutic strategies to enhance aged MuSC function in regeneration. Here we capitalize on multiparametric single-cell mass cytometry(Bjornson et al., 2013; Porpiglia et al., 2017; Spitzer and Nolan, 2016), which allows discovery of novel cell subsets within rare stem cell populations, to determine whether alterations in the relative abundance of MuSC subsets or in the regulation of their signaling networks is responsible for the decline of skeletal MuSC function in the course of aging. High dimensional single-cell analysis of MuSCs enabled us to identify two functionally and molecularly distinct subsets, defined by differential cell surface expression of CD47. CD47^lo^ MuSCs retain stem cell function, whereas CD47^hi^ MuSCs are defective in self-renewal. With aging, a shift in polyadenylation site choice leads to a marked increase in the relative abundance of CD47^hi^ MuSCs with impaired regenerative function.

CD47, known as the “don’t eat me signal” on cancer cells(Jaiswal et al., 2009), has only recently been implicated in myogenesis. A recent study revealed that CD47 signaling promotes proliferation of young MuSCs in the context of hypertrophy modeled by mechanical stress, leading to myonuclear accretion(Kaneshige et al., 2022). Here we uncover a new role for CD47 in MuSCs, in the context of aging and regeneration. We show that in contrast to young MuSCs, CD47 is markedly increased on the cell surface of aged MuSCs due to alternative polyadenylation resulting from increased U1 snRNA expression. Further, our study uncovers an unexpected age-dependent role for CD47 signaling in a paracrine loop that inhibits proliferation of the residual functional CD47^lo^ aged MuSC subset and has a pleiotropic negative effect on regenerative function in aged muscle. To overcome this age-related dysfunction, we identify two means to surmount aberrant CD47 signaling, via antisense morpholino oligonucleotides or immune blockade, which lead to enhanced regeneration. These findings have broad implications for sarcopenia and provide fresh insights into aged stem cell dysfunction of broad relevance to regenerative medicine.

## RESULTS

### CD47 expression levels distinguish functionally and molecularly distinct aged muscle stem cell subsets

We sought to identify cell surface markers that could distinguish novel MuSC subsets whose relative proportion is altered in the course of aging. We interrogated our previously described cell surface marker screen of single Pax7-ZsGreen muscle cells(Porpiglia et al., 2017) and found that CD47 distinguished two Pax7-ZsGreen^+^ MuSC subsets, CD47^lo^ and CD47^hi^ (Fig. 1A). To characterize these MuSC subsets in greater depth we performed CyTOF analysis. We added a CD47 antibody to our previously established CyTOF panel, which contained antibodies against surface markers CD9 and CD104, to distinguish MuSCs from progenitors P1 and P2, as well as CD44 and CD98, to identify activated MuSCs(Porpiglia et al., 2017). Analysis of the high-dimensional CyTOF data, using X-shift clustering paired with single-cell force directed layout visualization, revealed three distinct clusters within the *α*_7_integrin^+^/CD34^+^ MuSC population, distinguished by differential expression of CD47 and the transcription factor Pax7 (Fig. S1A, Fig.1B). The most prominent cluster was Pax7^hi^/CD47^lo^ (Fig. 1B, left), and the remaining two clusters were Pax7^lo^/CD47^int-hi^ (Fig. 1B, right). Further analysis of these subsets by CyTOF revealed that in addition to expressing low levels of Pax7 the CD47^hi^ subset expressed higher levels of the transcription factor Myf5, which marks activated stem cells, compared to the CD47^lo^ subset. Moreover, the expression of the myogenic transcription factors MyoD and Myogenin, characteristic of committed cells, was low in both MuSC subsets, as expected (Fig. S1B).

**Fig. 1.**
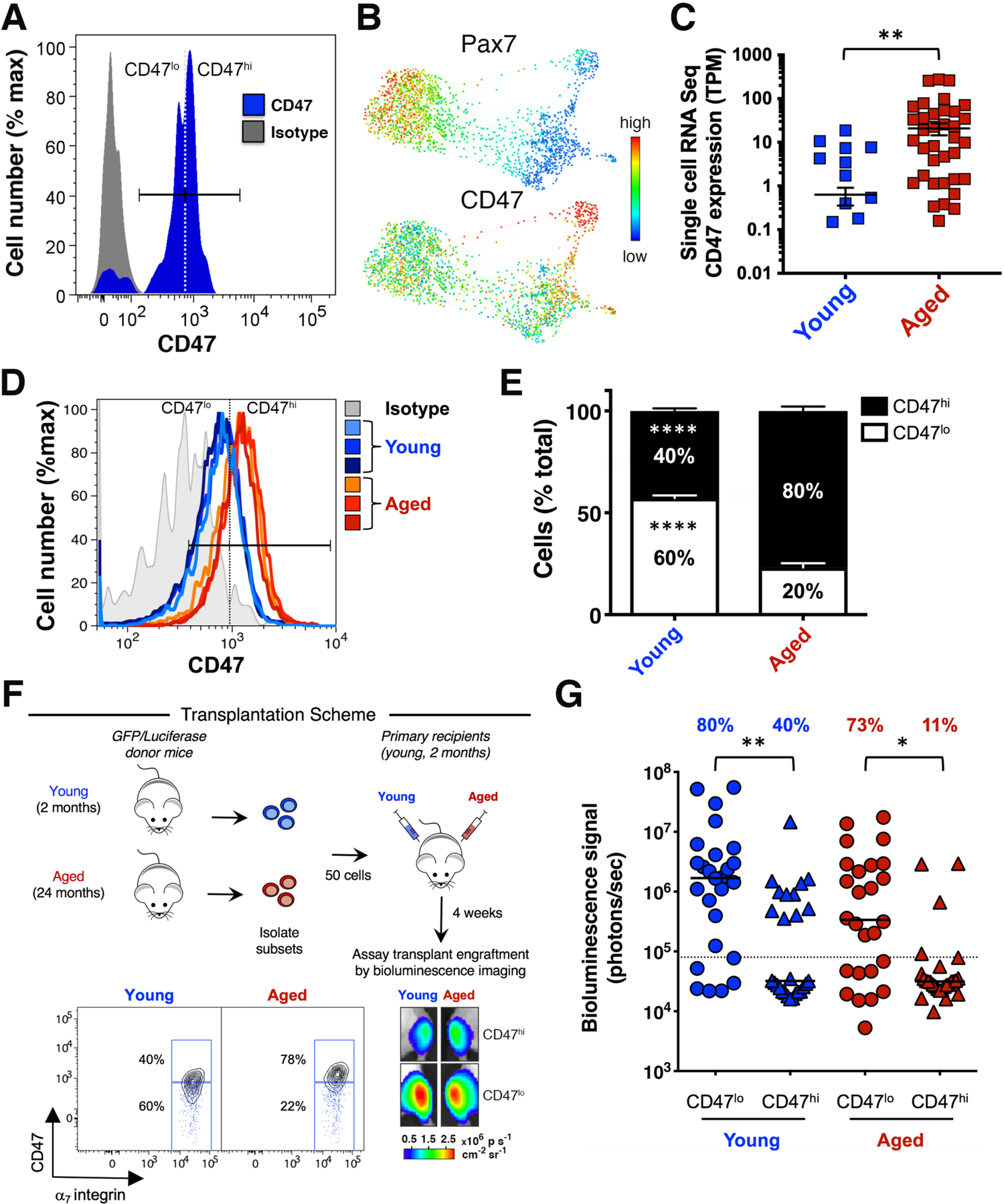
CD47 expression levels distinguish functionally and molecularly distinct aged muscle stem cell subsets. **(A)** Cell surface marker screening panel analysis of muscle stem cells (MuSCs). A single-cell suspension of *Tibialis Anterior* (TA) and *Gastrocnemius* (GA) muscle isolated from Pax7-ZsGreen reporter mice was stained with 176 cell surface antibodies and analyzed by fluorescence-based flow cytometry as previously described(Porpiglia et al., 2017). MuSCs are identified as ZsGreen^+^ cells. The histogram overlay shows CD47 expression in ZsGreen^+^ cells. The filled gray histogram represents the CD47 isotype control. **(B)** CyTOF mass cytometry workflow. TA and GA muscles from young mice were triturated, digested to a single-cell suspension, stained with isotope-chelated antibodies and run through the CyTOF instrument. Stained cells were passed through an inductively coupled plasma, atomized, ionized, and the elemental composition was mass measured. Signals corresponding to each elemental tag were correlated to the presence of the respective isotopic marker. Gated Live/Lineage^-^/*α*_7_integrin^+^/CD34^+^ MuSCs were analyzed with the X-shift algorithm (K=30 was auto-selected by the switch-point finding algorithm) yielding 3 clusters that were visualized using single-cell force-directed layout. Up to 2000 cells were randomly selected from each X-shift cluster, each cell was connected to 30 nearest neighbors in the phenotypic space and the graph layout was generated using the ForceAltas2 algorithm as previously described(Porpiglia et al., 2017). Expression levels of the myogenic transcription factor Pax7 (upper graph) and the surface marker CD47 (lower graph) were visualized (representative experiment, n= 3 mice; 4 independent experiments). **(C)** Scatter plot shows CD47 expression in young (2 months) and aged (24 months) MuSCs measured by single-cell RNA-seq analysis (mean ± SEM). Two-tailed unpaired t-test analysis with Welch’s correction was used to determine the difference in CD47 expression between young and aged MuSCs. **(D)** CD47 protein expression in young and aged MuSCs measured by flow cytometry. TA and GA muscles were triturated, digested to a single-cell suspension, stained with fluorophore-conjugated antibodies to lineage markers (CD45, CD31, CD11b, Sca1), *α*_7_integrin, CD34, CD47 and analyzed by fluorescence-based flow cytometry. MuSCs were identified as Live/Lin^-^/*α*_7_integrin^+^**/**CD34^+^ cells. The histogram overlay shows CD47 expression in Live/Lin^-^/*α*_7_integrin^+^**/**CD34^+^ MuSCs, from young (blue histograms) and aged (red histograms) mice (representative experiment, n= 3 young mice, 3 aged mice). The filled gray histogram represents the FMO + CD47 isotype control. **(E)** Quantification of the relative abundance of MuSC subsets. Stacked columns indicate the relative proportion of each subset (CD47^lo^ in white, CD47^hi^ in black) within the Live/Lin^-^/*α*_7_integrin^+^**/**CD34^+^ MuSC population in muscle isolated from young and aged mice (mean ± SEM from n=9 mice, 3 independent experiments). Two-way ANOVA analysis with Sidak correction for multiple comparisons was used to determine difference in the abundance of the individual subsets between young and aged MuSCs. **(F)** Scheme depicting the *in vivo* assay of regenerative capacity. Hindlimb muscles isolated from young and aged GFP^+^/Luciferase^+^ mice were digested to a single-cell suspension. CD47^hi^ and CD47^lo^ MuSC (Lin^-^/*α*_7_integrin^+^**/**CD34^+^) subsets were identified, as shown in the biaxial plot of CD47 (y axis) by *α*_7_integrin (x axis), sorted and transplanted (50 cells/injection) into the TA muscle of hindlimb-irradiated NOD/SCID mice. Overlaid blue dots in the biaxial plot indicate the FMO + CD47 isotype control. Representative BLI images at 4 weeks post-transplant are shown (lower right panel). **(G)** Scatter plot shows the percentage of transplants from each condition that engrafted above threshold (dashed line, 80,000 photons/s) into recipient tissue and the BLI signal intensity (y axis). Line represents the median BLI signal (n= 26 mice, 3 independent experiments). Kruskal Wallis test with significance determined by Dunn’s multiple comparisons test was performed to determine the engraftment difference between CD47^lo^ and CD47^hi^ MuSC subset isolated from young or aged mice. *, ** and **** represent statistical significance at p*≤*0.05, p*≤*0.01 and p*≤*0.0001 respectively.

Single-cell RNA-seq analysis of young and aged *α*_7_integrin^+^/CD34^+^ MuSCs revealed a marked increase in CD47 mRNA expression during aging (Fig. 1C), which was confirmed at the protein level by flow cytometry analysis (Fig. 1D). Specifically, we found that the proportion of CD47^hi^ MuSCs, as well as the CD47 signal intensity, increased with aging (Fig. 1D, E; Fig. S1C). These data demonstrate the presence of two previously unrecognized subsets within the MuSC population, CD47^lo^ and CD47^hi^, whose relative abundance shifts during aging, leading to accumulation of a CD47^hi^ subset.

To assess their regenerative potential, CD47^lo^ and CD47^hi^ *α*_7_integrin^+^/CD34^+^ MuSC subsets were isolated from young and aged GFP/Luciferase mice and transplanted into the irradiated *Tibialis Anterior* (TA) muscles of NOD/SCID mice. At 4 weeks post-transplant, the contribution of the donor cells to regenerated damaged tissues was determined by bioluminescence imaging (BLI) (Fig. 1F), as previously described(Sacco et al., 2008). Strikingly, based on the engrafted transplants, the CD47^lo^ subset isolated from young and aged donor mice exhibited the highest regenerative potential, as evidenced by high engraftment frequencies and BLI signal intensity (Fig. 1F, G). We evaluated stem cell repopulation in primary recipients by flow cytometric detection of donor-derived GFP^+^ MuSCs as a fraction of the total recipient MuSC population (Lin^-^/α_7_ integrin^+^/CD34^+^), 4 weeks after transplantation. We found that CD47^lo^ MuSCs from both young and aged donors gave rise to a larger GFP^+^ fraction of the total MuSC population in primary recipients than did CD47^hi^ MuSCs (Fig. S1D). These findings suggest that elevated CD47 may play a role in the reduced engraftment seen in the unfractionated aged MuSC population(Cosgrove et al., 2014).

### Alternative polyadenylation regulates CD47 expression at the onset of myogenic differentiation and is altered in aged muscle stem cells

In order to resolve the mechanism underlying the high expression of CD47 in aged MuSCs, we investigated the post-transcriptional regulation of CD47 expression. CD47 has been reported to undergo alternative polyadenylation in human cell lines leading to differential subcellular localization(Berkovits and Mayr, 2015; Ma and Mayr, 2018). The short 3’ UTR generates a CD47 protein that traffics primarily to the endoplasmic reticulum, whereas the long 3’ UTR, which binds a complex containing the RNA binding protein HuR, generates a protein that localizes to the cell surface(Berkovits and Mayr, 2015). HuR has been shown to increase at the onset of myogenic differentiation(Apponi et al., 2011; Figueroa et al., 2003; van der Giessen and Gallouzi, 2007). Hence, we hypothesized that the long 3’UTR isoform of the CD47 transcript was preferentially expressed during myogenic differentiation, resulting in increased CD47 protein localization at the cell surface. This hypothesis could explain the role of differential expression of CD47 protein on the surface of MuSC subsets in young and aged mice.

To assess whether stem and progenitor cells exhibited differential localization of CD47 protein, we measured its expression on both the cell surface and intracellularly by flow cytometry. We performed surface staining followed by cell permeabilization and intracellular staining using the same CD47 antibody labeled with different fluorophores. In MuSCs from young mice, CD47 expression was predominantly intracellular (Fig. 2A), whereas in progenitors from the same mice CD47 expression was predominantly on the cell surface (Fig. S2A). In aged mice, the relative abundance of MuSCs expressing surface CD47 and levels of surface CD47 protein were significantly greater than in young mice (Fig. 2A-C). However, these differences were not observed in progenitor cells from young and aged mice (Fig. S2A, B). These results show that CD47 localization on the cell surface, which increases with myogenic progression, increases prematurely in aged MuSCs.

**Fig. 2.**
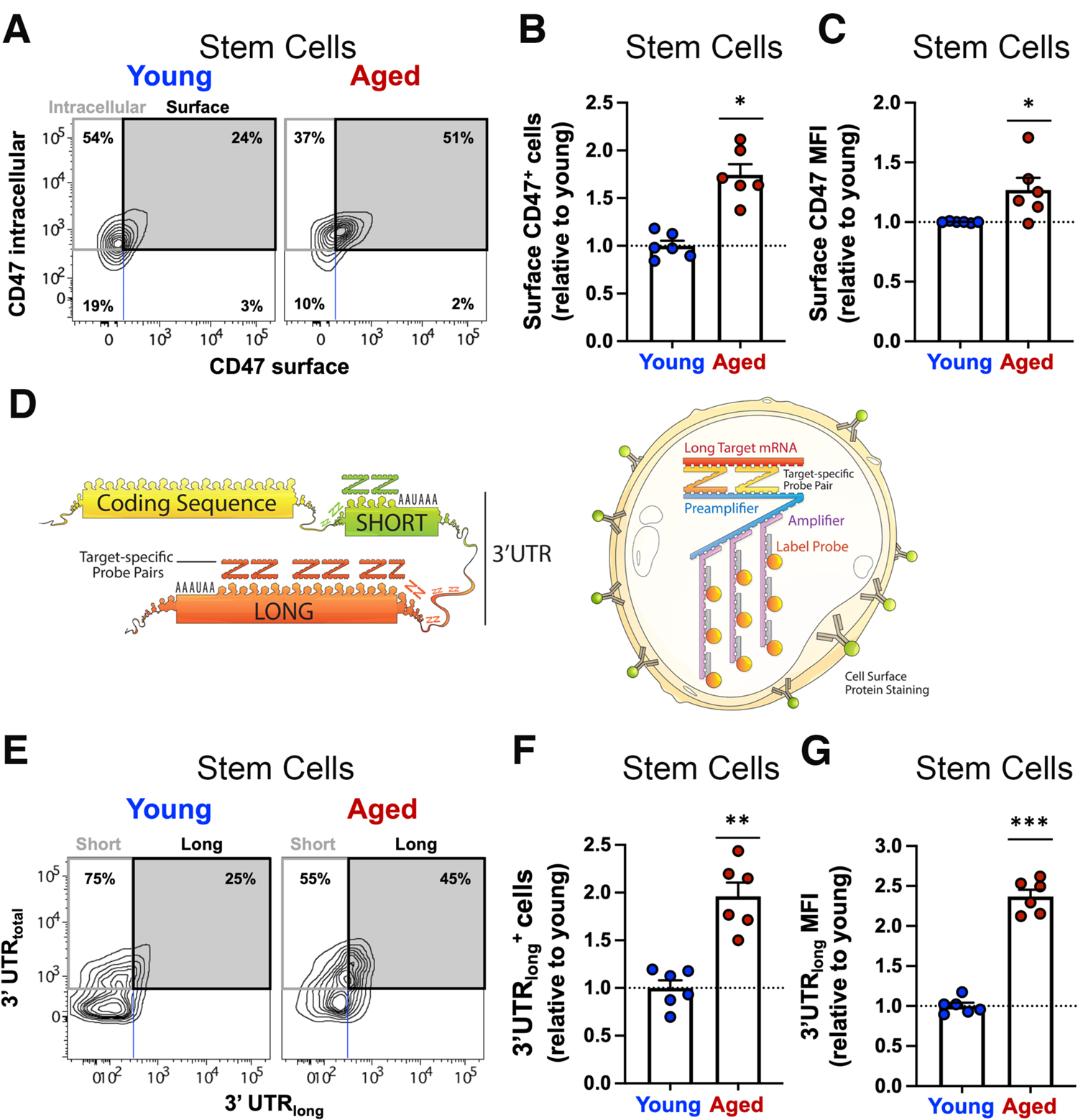
Alternative polyadenylation regulates CD47 expression at the onset of myogenic differentiation and is altered in aged muscle stem cells. **(A)** TA and GA muscles were isolated from young (2 months) and aged (24 months) mice, and digested to a single cell suspension that was stained using antibodies against lineage markers (CD45, CD11b, CD31, Sca1)-APCCy7, *α*_7_integrin-PE, CD9-APC and CD47-BV605. Cells were then fixed, permeabilized and stained for CD47 intracellularly using a different conjugate, CD47-PECy7. A biaxial dot plot of intracellular CD47-PECy7 (y axis) by surface CD47-BV605 (x axis) depicts the distribution of CD47 expression on the surface or intracellularly in stem cells from young (left panels) and aged (right panels) mice (representative sample). **(B)** Surface CD47^+^ cells in (A) are quantified. The bar graph indicates the abundance of aged surface CD47^+^ stem cells relative to young (mean ± SEM, n=6 mice, 2 independent experiments). Two-way ANOVA analysis was used to determine the difference between young and aged populations. **(C)** The bar graph shows the expression level of CD47 protein on the surface of aged MuSCs (*α*_7_integrin^+^/CD34^+^), measured by flow cytometry as median fluorescence intensity (MFI) (mean ± SEM from n=6 mice, 2 independent experiments) and quantified relative to young MuSCs. Two-tailed paired t-test analysis was used to determine the difference in surface CD47 MFI between young and aged stem cells. **(D)** Scheme depicting the CD47 coding sequence followed by the 3’ untranslated region (UTR). The 3’ UTR contains two functional polyadenylation sites (PAS): alternative polyA site selection generates messenger RNA (mRNA) transcripts of different length. RNA in situ hybridization probes were custom designed to differentiate between the short and long 3’UTR isoforms of CD47 mRNA using the PrimeFlow RNA assay. About 8000 fluorophores labeled each target mRNA. **(E)** TA and GA muscles were isolated from young and aged mice, and digested to a single cell suspension that was stained using antibodies against lineage markers (CD45, CD11b, CD31, Sca1)-APCCy7, *α*7 integrin-PE, CD9-APC and CD47-BV605. Cells were then fixed, permeabilized and stained for CD47 intracellularly using a different conjugate CD47-PECy7. Finally, cells were stained for the different CD47 mRNA isoforms using the custom-designed probes targeting the total or long 3’UTR isoform of CD47 mRNA and the PrimeFlow RNA assay kit according to manufacturer’s instructions. The total and long 3’UTR isoform of CD47 mRNA were labeled with AF647 and AF750 respectively. A biaxial dot plot of CD47 mRNA-3’UTR_total_ (y axis) by CD47 mRNA-3’UTR_long_ (x axis) depicts the distribution of cells expressing the short isoform (upper left quadrant) or the long isoform (upper right quadrant) as a fraction of total CD47 mRNA (upper left and right quadrants combined) in stem cells from young (left panels) and aged (right panels) mice (representative sample). **(F)** Bar graph indicates the relative increase in the abundance of aged stem cells expressing the CD47 mRNA with long 3’UTR compared to young stem cells (mean ± SEM, n=6 mice, 2 independent experiments). Two-tailed paired t-test analysis was used to determine difference in the proportion of stem cells expressing the CD47 mRNA with long 3’UTR between young and aged samples. **(G)** Bar graph shows the expression level of the CD47 mRNA with long 3’UTR in aged stem cells, measured by flow cytometry as median fluorescence intensity (MFI) and quantified relative to young MuSCs (mean ± SEM, n=6 mice, 2 independent experiments). Two-tailed paired t-test analysis was used to determine difference in the expression level of CD47 mRNA with long 3’UTR between young and aged stem cells. *, ** and *** represent statistical significance at p*≤*0.05, p*≤*0.01 and p*≤*0.001 respectively.

To map and determine the abundance of the different CD47 mRNA 3’UTR isoforms, we performed transcript alignment from different species in combination with analysis of published datasets obtained by 3’ region extraction and deep sequencing of murine cell lines(Hoque et al., 2013) (Fig. S2C, D). We identified three previously unrecognized murine CD47 mRNA isoforms with 3’UTRs of different lengths that showed, from shortest to longest, progressively lower usage (Fig. S2C, D). We focused our analysis on the most prevalent isoforms containing the polyadenylation sites, PAS1 or PAS2, herein referred to as short and long, respectively (Fig. 2D, left panel).

To measure the distribution of short and long 3’UTR transcripts simultaneously in stem and progenitor cells, we used branched DNA technology(van Buuren et al., 2018; Hensel et al., 2021) to specifically label individual CD47 mRNA isoforms in single-cells with target-specific probes spanning 600-800 bases upstream of PAS1 and PAS2 (Fig. 2D, right panel). We stained single-cell suspensions of muscle cells from either young or aged mice, first with antibodies against the surface markers *α*_7_ integrin and CD9, to distinguish both stem (Lin^-^/*α*_7_ integrin^+^/CD9^int^) and progenitor (Lin^-^/*α*_7_ integrin^+^/CD9^hi^) cells (Fig. S2E), and then intracellularly with probes specific to the different isoforms of the CD47 mRNA (Fig. 2D). The progenitor cell gate included the two previously described P1 and P2 progenitor cell populations(Porpiglia et al., 2017).

In aged mice, the proportion of cells and levels of expression of the long CD47 mRNA isoform were comparable in both stem and progenitor cells (Fig. 2E, and Fig. S2F). This contrasted with young mice where more progenitor cells than MuSCs expressed the long CD47 mRNA isoform (Fig. 2E, and Fig. S2F). Furthermore, the relative abundance of MuSCs expressing the long CD47mRNA isoform and the expression level of the long CD47 mRNA isoform were higher in aged compared to young stem cells (Fig. 2F, G). We validated these results using an orthogonal approach, isoform specific q-RT-PCR (Fig. S2H). These data indicate alternative polyadenylation as the mechanism underlying increased surface CD47 expression.

### U1 snRNA skews the balance between CD47 mRNA isoforms in aged muscle stem cells

To gain mechanistic insight into CD47 mRNA alternative polyadenylation in aged MuSCs, we aligned transcripts from different species, with a focus on the region upstream of the proximal polyadenylation site (PAS1). This analysis identified a conserved binding site for U1 small nuclear RNA (U1 snRNA), an established component of the RNA splicing machinery(Nilsen, 2003; Wahl et al., 2009) (Fig. 3A). Critically, U1 snRNA also plays a role in alternative polyadenylation facilitating proximal polyadenylation site readthrough(Berg et al., 2012; Kaida et al., 2010; de Morree et al., 2019). We hypothesized that differential expression of U1 snRNA might be responsible for the increased abundance of the long CD47 mRNA isoform in aged MuSCs. We measured U1 snRNA expression in sorted young and aged MuSCs and found that its expression was significantly increased in aged compared to young MuSCs (Fig. 3B). By contrast, no significant differences were observed for other factors implicated in alternative polyadenylation(Tian and Manley, 2016) (Fig. S3A).

**Fig. 3.**
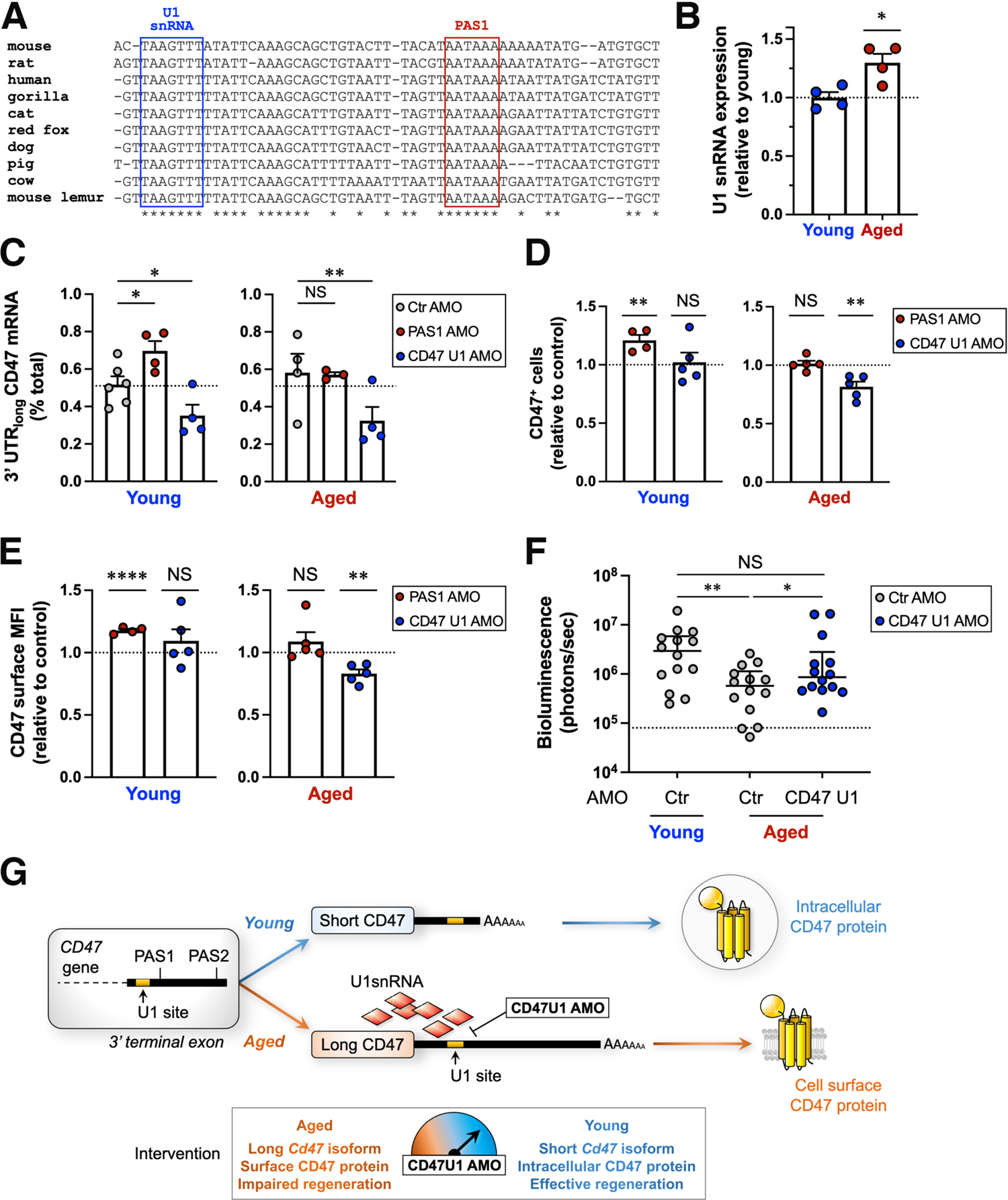
U1 snRNA skews the balance toward short 3’UTR CD47 mRNA isoforms in aged muscle stem cells. **(A)** The CD47 mRNA sequence upstream of polyadenylation site 1 (PAS1) from mouse, human, gorilla, cat, red fox, dog, pig, cow and mouse lemur were aligned using MAFFT online. A highly conserved U1 snRNA binding site (blue box) was identified upstream of PAS1 (red box). **(B)** Scatter plot shows the expression levels of U1 snRNA in sorted aged MuSCs relative to young MuSCs, measured by RT-qPCR (mean ± SEM, n= 4 young and 4 aged mice, 2 independent experiments). Two-tailed paired t test was performed to determine the difference in U1 snRNA expression levels between young and aged MuSCs. **(C)** Bar graph shows the relative abundance of CD47 mRNA with long 3’ UTR in young and aged MuSCs treated in culture for 24 hours with antisense morpholino oligonucleotides (AMO) to polyadenylation site 1 (PAS1, red), U1snRNA binding site on CD47 transcript (CD47 U1, blue), and control (ctr, gray) (mean ± SEM, 4 independent experiments). Two-way ANOVA analysis was used to determine difference in the abundance of CD47 mRNA with long 3’ UTR across the different conditions in young or aged sorted MuSCs. **(D, E)** Surface CD47 protein expression was measured by flow cytometry in young and aged MuSCs treated in culture for 32 hours with AMOs to PAS1, CD47 U1 and control AMO. **(D)** Bar graph shows the proportion of young (left) and aged (right) MuSCs expressing surface CD47 upon treatment with PAS1 (red) or CD47 U1 (blue) AMO relative to control AMO treated MuSCs (mean ± SEM, 3 independent experiments). Two-way ANOVA was performed to determine the difference in the fraction of surface CD47 expressing MuSCs between the different conditions in young or aged sorted MuSCs. **(E)** Bar graph shows the expression level of surface CD47 protein on young (left) and aged (right) MuSCs treated as described above, measured by flow cytometry as median fluorescence intensity (MFI) and quantified relative to control AMO treated MuSCs (mean ± SEM, 3 independent experiments). Two-way ANOVA was performed to determine the difference in surface CD47 expression levels between the different conditions in young, or aged sorted MuSCs. **(F)** Hindlimb muscles isolated from young and aged GFP^+^/Luciferase^+^ mice were digested to a single-cell suspension. MuSCs were sorted and treated overnight with AMOs to CD47 U1 (blue) or control (ctr) AMO (gray). Cells were then transplanted (100 cells/injection) into the TA muscle of hindlimb-irradiated NOD/SCID mice and mice were imaged weekly by BLI. Scatter plot shows the BLI signal intensity of the engrafted transplants (y axis), 3 weeks after transplantation. Line represents the median BLI signal with the interquartile range (n= 14 mice, 2 independent experiments). Kruskal Wallis test was performed to determine the engraftment difference between the different conditions. **(G)** Model for dysregulation of alternative polyadenylation in aged MuSCs and intervention with CD47U1 AMO. U1 snRNA is upregulated in aged MuSCs. U1 snRNA binds to its site on CD47 3’UTR upstream of the proximal polyadenylation site (PAS1), blocks usage of CD47 PAS1 site, and shifts the balance toward the long isoform, leading to increased surface CD47 expression. An AMO complementary to the U1 site on CD47 (CD47U1 AMO intervention) competes with U1 snRNA for its binding site and interferes with U1 snRNA function at the CD47 transcript. As a result, alternative polyadenylation is prevented, leading to increased production of the short CD47 isoform. This in turn results in decreased surface CD47 expression and improved MuSC engraftment to levels similar to those of young MuSCs. The speed dial shows that increasing levels of CD47U1 AMO shift the balance towards the short CD47 isoform and a more youthful molecular phenotype. *, ** and **** represent statistical significance at p*≤*0.05 p*≤*0.01, and p*≤*0.0001 respectively.

To test the role of U1 snRNA in the regulation of CD47 mRNA alternative polyadenylation, we designed an antisense morpholino oligonucleotide (AMO) complementary to the U1 snRNA binding site (CD47 U1 AMO), to block the binding of U1 snRNA to the CD47 mRNA 3’ UTR. In addition, we designed an AMO complementary to a conserved sequence in the PAS1 (PAS1 AMO), to compete with the polyadenylation complex and force the selection of a more distal PAS. We treated sorted young and aged MuSCs with the CD47 U1, PAS1 or control AMO and measured CD47 mRNA isoform abundance by RT-qPCR and surface CD47 protein expression by flow cytometry. Treatment with the PAS1 AMO led to an increase in the relative abundance of the long CD47 mRNA isoform and surface CD47 protein levels in young but not aged MuSCs, where the relative abundance of the long CD47 mRNA isoform and surface CD47 protein levels were already elevated at steady state (Fig. 3C-E). These data confirm our model that alternative polyadenylation is the mechanism underlying increased surface CD47 expression. By contrast, treatment with the CD47 U1 AMO led to a decrease in the relative abundance of the long CD47 mRNA isoform, with a consequent decrease in surface CD47 protein levels in aged MuSCs (Fig. 3C-E). These findings suggest that U1 snRNA controls CD47 alternative polyadenylation and that altered U1 snRNA expression contributes to the dysregulation of CD47 alternative polyadenylation in aged MuSCs. Consistent with alternative polyadenylation as a key regulatory mechanism for surface CD47 expression, we found that the complex (Elav1, Set, and Rac1) that controls CD47 localization is increased in aged MuSCs(Berkovits and Mayr, 2015) (Fig. S3B-D).

To determine the biological significance of reversing the relative abundance of the short and long CD47 mRNA isoforms in aged MuSCs, we performed transplantation assays with MuSCs treated with the CD47 U1 or control AMO. Strikingly, at 3 weeks post-transplantation, aged MuSCs treated with CD47 U1 AMO had significantly higher regenerative capacity compared to controls (Fig. 3F). Altogether, these data demonstrate that blocking U1 snRNA binding to CD47 mRNA in aged MuSCs rejuvenates their function in regeneration, by shifting the balance of CD47 mRNA isoforms toward the short isoform, and decreasing surface CD47 expression (Fig. 3G).

### Aberrant thrombospondin-1 signaling via CD47 inhibits the proliferative capacity of aged muscle stem cells

CD47 has dual functions as ligand and receptor(Oldenborg, 2013). Also known as the “don’t eat me signal” on neoplastic cells(Jaiswal et al., 2009; Majeti et al., 2009), CD47 is a ligand for SIRP-*α*, a receptor present on the surface of immune cells, which upon binding to CD47 delivers an inhibitory signal that blocks phagocytosis(Jaiswal et al., 2009). In addition, CD47 is known as Integrin Associated Protein (IAP) because it interacts with integrins(Brown and Frazier, 2001). Finally, CD47 also serves as a receptor for thrombospondin-1, a secreted cellular matrix protein that has been shown to inhibit the proliferation of endothelial cells(Feng et al., 2015; Gao et al., 2016; Kaur et al., 2013; Oldenborg, 2013). During aging, thrombospondin-1 has been shown to accumulate in several tissues, including skeletal and cardiac muscle(Isenberg and Roberts, 2020) and its expression in skeletal muscle has been shown to decrease in response to exercise(Hoier et al., 2013a, 2013b; Isenberg and Roberts, 2020; Kivelä et al., 2008). We found that *thrombospondin-1* transcripts are increased in the aged muscle stem cells compared to young (Fig. 4A). To establish whether increased *thrombospondin-1* expression could account for the reduced proliferative potential of aged MuSCs, we investigated the thrombospondin-1/CD47 signaling axis and performed a series of *in vitro* and *in vivo* experiments.

**Fig. 4.**
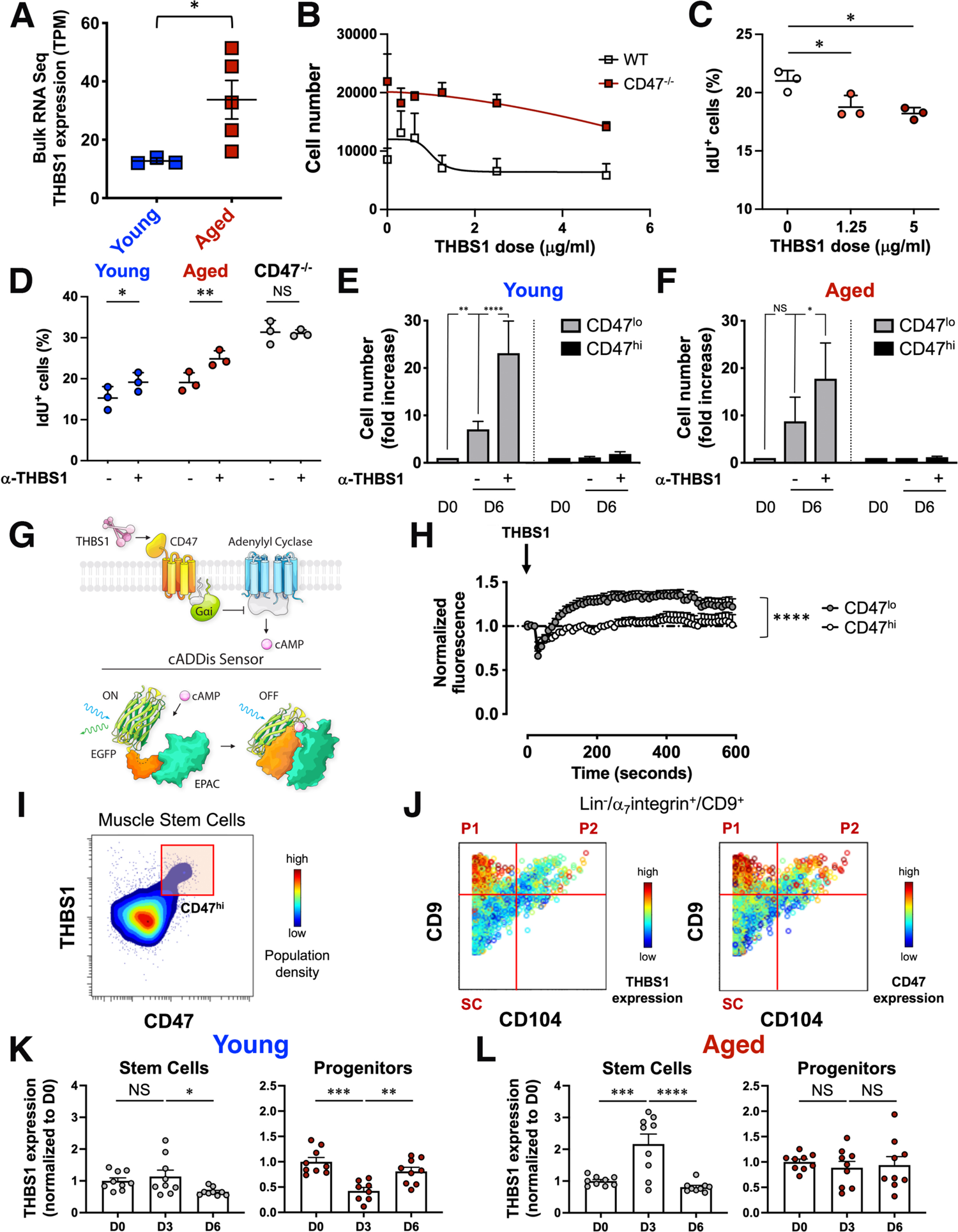
Aberrant thrombospondin-1 signaling via CD47 inhibits the proliferative capacity of aged muscle stem cells. **(A)** Scatter plot shows thrombospondin-1 (THBS1) expression in young and aged MuSCs measured by bulk RNA seq analysis (mean ± SEM, n=3 young mice, n=5 aged mice). Two-tailed unpaired t-test analysis with Welch’s correction was used to determine the difference in THBS1 expression between young and aged MuSCs. **(B)** Sorted *α*_7_integrin^+^/CD34^+^ MuSCs from young wild type and CD47^-/-^ mice were cultured for one week in the presence of increasing doses of THBS1 (0.6μg/ml-5μg/ml) and proliferation was quantified by cell count (mean ± SEM, n=14 wild type replicates, n=4 CD47^-/-^ replicates, 4 independent experiments). **(C)** Scatter plot shows the fraction of IdU^+^ MuSCs measured by CyTOF analysis in response to a 6-day treatment of young MuSCs with increasing doses of recombinant THBS1 (mean ± SD, n= 3 young mice). One-way ANOVA analysis with Tukey’s correction for multiple comparisons was used to determine difference in the fraction of IdU^+^ cells between untreated and THBS1 treated young MuSCs. **(D)** Scatter plot shows the fraction of IdU^+^ MuSCs measured by CyTOF analysis in response to a 6-day treatment of sorted wild type (young, aged) and CD47^-/-^ aged MuSCs with a blocking antibody to THBS1 (*α*-THBS1) (mean ± SD, n= 3 mice per genotype). Two-way ANOVA with Sidak’s correction for multiple comparisons was performed to determine the difference in the fraction of IdU^+^ cells between *α*-THBS1 treated and control sorted MuSCs from young, aged or CD47^-/-^ mice. **(E, F)** Quantification of MuSC subset expansion after THBS1 blockade. The CD47^lo^ and CD47^hi^ MuSCs subsets were sorted from young **(E)** and aged **(F)** mice, cultured in growth media for 6 days on biomimetic hydrogels in the presence (+) or absence (-) of a blocking antibody to THBS1, and cell number was quantified by cell count at day 0 (D0) or at day 6 (D6) after culture. Bar graph shows cell number normalized to D0 for each subset (mean ± SEM from n= 7 young replicates, n=3 aged replicates). Two-way ANOVA analysis with Tukey’s correction for multiple comparisons was used to determine difference between conditions in CD47^lo^ and CD47^hi^ subsets isolated from young (E) or aged (F) mice. **(G)** (Top scheme) THBS1–CD47–cAMP signaling axis. THBS1–CD47 signals through Guanine nucleotide-binding protein G_i_’s alpha subunit which inhibits adenylyl cyclases to reduce cAMP levels. (Bottom scheme) The cADDis downward sensor is a fluorescent cAMP sensor capable of detecting changes in cAMP concentration in living cells. cAMP binding to the cADDis downward sensor reduces GFP fluorescence. **(H)** CD47^lo^ and CD47^hi^ MuSCs were sorted from young mice and transfected overnight with a baculovirus encoding the cAMP sensor. The following day MuSC subsets were treated with THBS1 and individually imaged with a confocal microscope at 10 sec intervals for 5 minutes. The graph shows the average normalized GFP fluorescence of CD47^lo^ (gray) and CD47^hi^ (white) MuSCs over time (mean ± SEM, 4 mice, 2 independent experiments). Two-tailed paired t-test analysis was used to determine the difference in normalized fluorescence expression between CD47^lo^ and CD47^hi^ MuSC subsets after stimulation with THBS1. **(I)** CyTOF analysis of young hindlimb muscle showing *α*_7_integrin^+^**/**CD34^+^ MuSCs from young mice. Representative biaxial dot plots of THBS1 (y axis) by CD47 (x axis) colored by population density shows a CD47^hi^ MuSC subset that expresses high levels of THBS1. **(J)** CyTOF analysis of young hindlimb muscle showing the entire myogenic compartment defined as Live/Lin^-^**/***α*_7_ integrin^+^**/**CD9^+^ cells. Representative biaxial dot plots of CD9 (y axis) by CD104 (x axis) colored by THBS1 expression (left) and CD47 expression (right) highlights that the progenitor population P1 expresses the highest level of both THBS1 and CD47. **(K, L)** Intracellular THBS1 protein measurement by CyTOF during regeneration. Mice were acutely injured by notexin injection in the TA and GA muscles, 6 or 3 days prior to tissue collection. Muscle tissues of the indicated groups were simultaneously collected at day 0, stained with isotope-chelated antibodies, run through the CyTOF instrument and analyzed. Bar graphs show THBS1 protein levels, normalized to day 0 (D0), in SC (left) and P1 (right) population from young (K) and aged (L) mice during an injury time course (mean ± SEM, n=9 mice from 2 independent experiments). One-way ANOVA analysis with Tukey’s correction for multiple comparisons was used to determine the difference in THBS1 expression between the indicated timepoints in SC or P1 cells from young (K) or aged (L) mice. *, **, *** and **** represent statistical significance at p*≤*0.05, p*≤*0.01, p*≤*0.001 and p*≤*0.0001 respectively.

First, to determine whether exposure to thrombospondin-1 suppresses MuSC proliferation *in vitro*, MuSCs from wild type and CD47^-/-^ mice were cultured for one week in the presence of increasing concentrations of recombinant thrombospondin-1. As the dose of thrombospondin-1 increased, the number of wild type MuSCs significantly decreased, demonstrating that thrombospondin-1 suppresses MuSC proliferation *in vitro* (Fig. 4B). Furthermore, decreased incorporation of iododeoxyuridine (IdU), decreased numbers of Ki67^+^ cells and <0.5% of cells expressing cleaved PARP confirmed that the thrombospondin-1-mediated decrease in cell number was due to inhibition of cell proliferation rather than increased apoptosis (Fig. 4C, Fig. S4A, B). Notably, MuSCs from CD47^-/-^ mice, which exhibited constitutively high proliferation rates, did not respond to thrombospondin-1 treatment, confirming that thrombospondin-1-mediated suppression of proliferation depends on CD47 (Fig. 4B). To determine the mechanism of thrombospondin-1-mediated inhibition of proliferation we measured levels of cell cycle inhibitors (CDK interacting protein/Kinase inhibitor protein (Cip/Kip CKI)) after thrombospondin-1 treatment *in vitro*, based on previous findings(Gao et al., 2016) (Fig. S4C, D). q-RT-PCR analysis of young MuSCs revealed that transcripts for p21 (CDKN1a), p27(CDKN1b) and p57 (CDKN1c) increased significantly after thrombospondin-1 treatment (Fig. S4D), and CyTOF analysis confirmed the increase in p57 at the protein level (Fig. S4C).

### Thrombospondin-1 blockade restores the proliferative capacity of aged muscle stem cells *in vitro*

To assess whether thrombospondin-1 blockade could mitigate the proliferative defect of aged MuSCs *in vitro,* we treated them with a thrombospondin-1 antibody that blocked its interaction with CD47(Annis et al., 2006). Proliferation of aged MuSCs was restored to levels comparable to untreated young MuSCs, whereas control CD47^-/-^ MuSCs were refractory to this treatment, underscoring the specificity of the blocking antibody (Fig. S4E, F). Finally, an *in vitro* IdU incorporation assay followed by CyTOF analysis confirmed that the increase in cell number upon thrombospondin-1 blockade, was due to an increase in cell proliferation, as shown by an increase in the fraction of IdU^+^ cells in both young and aged MuSCs but not in CD47^-/-^ MuSCs (Fig. 4D). To establish the cellular target of thrombospondin-1, we sorted CD47^lo^ and CD47^hi^ MuSC subsets from young and aged mice and treated them *in vitro* with the blocking antibody to thrombospondin-1 for one week. Strikingly, in response to the blocking antibody, proliferation increased in the CD47^lo^ MuSC subset isolated from young and aged mice, while no significant changes were observed for the CD47^hi^ subset (Fig. 4E, F). These findings demonstrate that thrombospondin-1 preferentially suppresses the proliferation of MuSCs expressing low levels of CD47 *in vitro*. These data suggest that MuSC subsets that express different levels of CD47 exhibit different sensitivities to thrombospondin-1, with consequences for recruitment of intracellular signaling pathways, cell cycle phase and cell fate determination.

To determine whether the CD47^lo^ and CD47^hi^ MuSC subsets differed in their cell cycle status, we performed CyTOF analysis of MuSCs harvested during an *in vivo* time course of recovery from acute injury. Upon injury CD47^lo^/Pax7^hi^ MuSCs incorporated high levels of IdU demonstrating increased proliferation, which was reflected in the increased relative abundance (Fig. S4G, H). By contrast, the CD47^hi^/Pax7^lo^ MuSCs incorporated little IdU and their relative abundance decreased during the course of injury (Fig. S4G, H). These findings provide additional evidence that CD47^lo^ MuSCs represent self-renewing stem cells.

To determine differential signaling responses of MuSCs to thrombospondin-1 stimulation, we focused on cAMP signaling, a key regulator of cell proliferation(Dumaz and Marais, 2005; Stork and Schmitt, 2002), as thrombospondin-1 decreases cAMP levels through activation of G_i_ proteins(Wang et al., 1999; Yao et al., 2011). Basal cAMP levels were significantly higher in sorted CD47^lo^ MuSCs compared to CD47^hi^ MuSCs (Fig. S4I). To determine whether CD47^lo^ and CD47^hi^ MuSC subsets differed in their ability to modulate cAMP signaling in response to thrombospondin-1 stimulation, we employed live single-cell confocal imaging of sorted MuSC subsets expressing a fluorescent cAMP downward sensor(Tewson et al., 2016) (Fig. 4G). Using this assay, an increased GFP signal is a readout for decreased cAMP levels (Fig. 4G). CD47^lo^ and CD47^hi^ MuSCs were isolated from young mice and transfected with a baculovirus encoding the cAMP downward sensor. MuSC subsets were then treated with thrombospondin-1 and individually imaged every 10 sec for 5 minutes. Upon thrombospondin-1 treatment, the GFP signal increased in the CD47^lo^ but not in the CD47^hi^ MuSC subset (Fig. 4H), demonstrating that thrombospondin-1 signals through cAMP in the CD47^lo^ but not in the CD47^hi^ MuSC subset. The phenotypic analysis of the MuSC subsets, together with the cAMP signaling studies, indicate that the CD47^lo^ and CD47^hi^ MuSCs subsets are both functionally and molecularly distinct.

To establish the cell type within skeletal muscle that produces and secretes thrombospondin-1, we capitalized on mass cytometry. Young and aged muscle cells were stained with antibodies against cell surface markers that distinguish myogenic stem and progenitor cells and an antibody against thrombospondin-1, which was detected intracellularly upon permeabilization. Data analysis showed that the CD47^hi^ MuSC subset expressed greater thrombospondin-1 levels than the CD47^lo^ MuSC subset (Fig. 4I).

We reasoned that thrombospondin-1 could have a physiological role during regeneration. Thus, we extended our analysis of thrombospondin-1 expression to cells within the Lin^-^/*α*_7_integrin^+^/CD9^+^ myogenic compartment, which includes, in addition to the stem cells (Lin^-^/*α*_7_integrin/CD9^int^), progenitor populations (Lin^-^/*α*_7_integrin/CD9^hi^) P1 and P2, distinguished by co-expression of CD9 and CD104, as we described previously(Porpiglia et al., 2017). Strikingly, P1 progenitor cells expressed significantly greater levels of both thrombospondin-1 and CD47 compared to MuSCs (Fig. 4J).

We hypothesized that during regeneration, after MuSCs expand, their P1 progeny secrete thrombospondin-1 creating a negative feedback loop that prevents stem cell exhaustion and promotes MuSC return to quiescence. To test this, we investigated the dynamics of thrombospondin-1 expression *in vivo*, in the context of recovery from acute injury. We performed an injury time course by notexin injection and measured thrombospondin-1 expression intracellularly in muscle stem and progenitor cells from young and aged mice by CyTOF. Thrombospondin-1 expression was low in young MuSCs compared to young P1 progenitor cells and further decreased at day 6 post injury (Fig. 4K, left panel). In young P1 progenitor cells thrombospondin-1 expression changed dynamically throughout the injury time course, decreasing significantly at day 3 post injury, a time during which MuSCs expand dramatically(Porpiglia et al., 2017), and increasing significantly at day 6 post injury (Fig. 4K, right panel). By contrast, thrombospondin-1 expression in the aged myogenic compartment exhibited a marked dysregulation. In aged MuSCs, thrombospondin-1 expression levels significantly increased at day 3 post injury, and decreased by day 6 (Fig. 4L, left panel). In aged P1 progenitor cells, thrombospondin-1 expression levels were low and did not change (Fig. 4L, right panel).

The dynamic change in thrombospondin-1 expression in young progenitor cells suggests a mechanism by which progenitor cell population density influences MuSC expansion. We propose that the population density of progenitor cells can be sensed by MuSCs through the secreted thrombospondin-1 that, when released by progenitor cells after MuSC expansion, acts in a paracrine fashion to suppress proliferation of the neighboring MuSCs and promote their return to quiescence.

### Thrombospondin-1 blockade *in vivo* activates aged muscle stem cells in the absence of injury

To test this model *in vivo*, we treated *Pax7^CreERT2^;Rosa26-LSL-Luc* mice, a transgenic mouse model in which endogenous MuSC numbers can monitored by bioluminescence imaging, with a thrombospondin-1 blocking antibody or IgG control regimen, which consisted of three intramuscular (i.m.) injections in the TA muscle at two-day intervals (Fig. 5A, scheme). Remarkably, *in vivo* treatment with the thrombospondin-1 blocking antibody was sufficient to significantly expand MuSCs in the absence of injury compared to the IgG control (Fig. 5A). These data suggest that thrombospondin-1 plays a role in preventing MuSC activation during homeostasis.

**Fig. 5.**
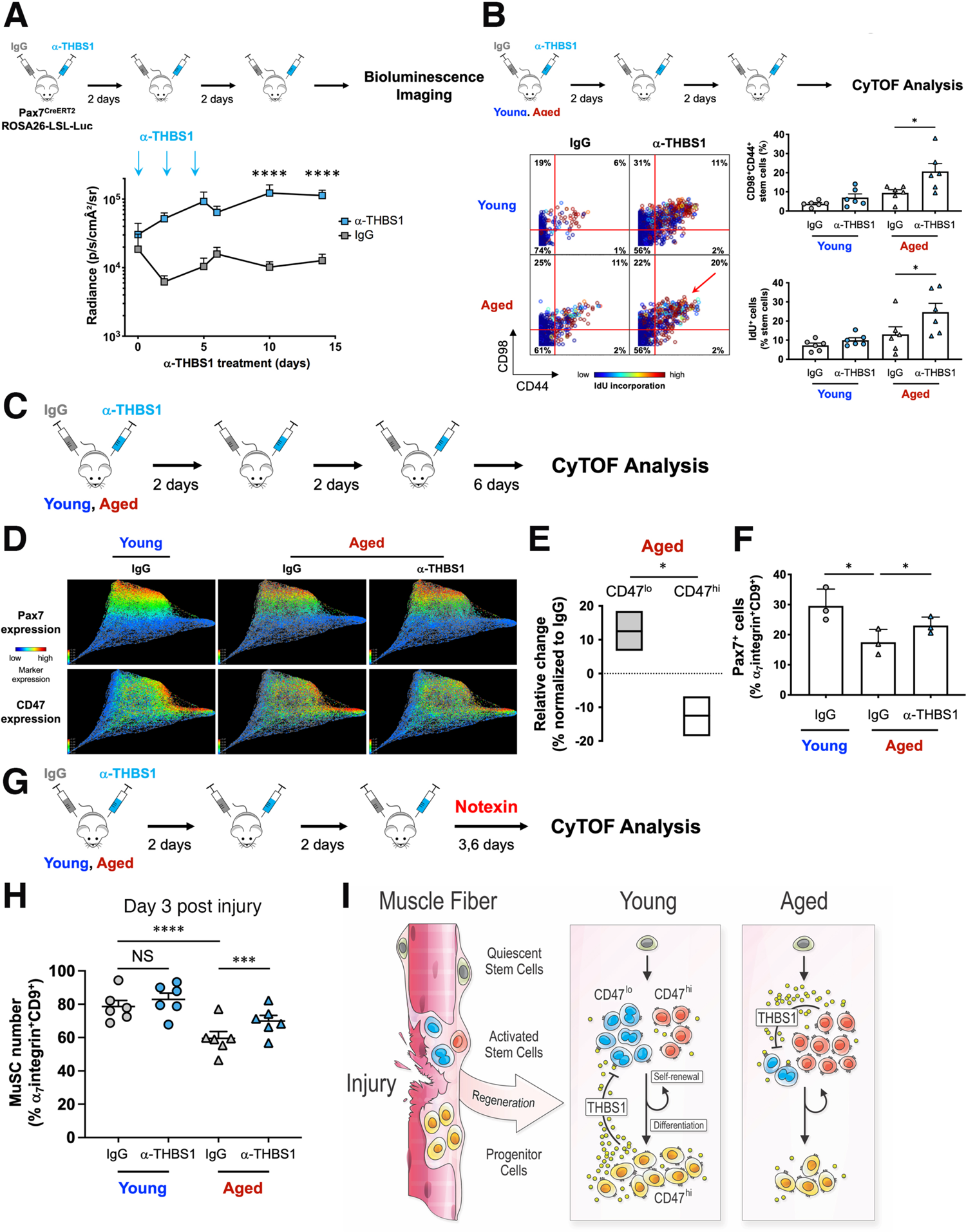
Thrombospondin-1 blockade *in vivo* activates aged muscle stem cells in absence of injury and enhances the regenerative response of aged muscle. **(A)** Experimental scheme (upper panel). Endogenous MuSC expansion was assayed by bioluminescence imaging (BLI) in Pax7^CreERT2^;Rosa26-LSL-Luc mice treated with tamoxifen (TAM). Mice were injected intramuscularly in the TA and GA muscles (three times, at two-days interval) with an antibody to THBS1 (*α*-THBS1) or a control IgG (contralateral leg) and imaged by BLI. The graph shows the summary of the BLI signal intensity (y axis) over time for each group (light blue, treatment with THBS1 blocking antibody; gray, treatment with IgG control) (mean ± SEM, n = 7 mice per condition, 2 independent experiments). Multiple t-tests with Holm-Sidak correction for multiple comparisons were used to determine the difference in bioluminescence signal between IgG control and *α*-THBS1 treated samples at the indicated time points. **(B)** Experimental scheme (upper panel). Young and aged mice were treated by intramuscular injections in the TA and GA muscles with a blocking antibody to THBS1 as in (A) and hindlimb muscle tissue was collected for CyTOF analysis at the end of treatment. (Left) Representative biaxial dot plots of CD98 (y axis) by CD44 (x axis) colored by channel show IdU incorporation in activated MuSCs, defined by co-expression of surface markers CD98 and CD44 (upper right quadrant). (Upper right) Bar graph indicates the proportion of CD98^+^/CD44^+^ activated MuSCs (mean ± SEM, n = 6 mice, 2 independent experiments). Two-tailed paired t-test was used to determine the difference in the abundance of CD98^+^/CD44^+^ subset between IgG control and *α*-THBS1 treated young or aged samples. (Lower right) Bar graph shows the proportion of IdU^+^ cells in the MuSC population (mean ± SEM, n = 6 mice, 2 independent experiments). Two-tailed paired t-test was used to determine the difference in the abundance of IdU^+^ stem cells between IgG control and *α*-THBS1 treated young or aged samples. **(C)** Experimental scheme. Young and aged mice were treated with a blocking antibody to THBS1 as in (B) and hindlimb muscle tissue was collected for CyTOF analysis, 6 days after the last THBS1 blocking antibody or IgG injection. **(D)** Live/Lineage^-^/*α*_7_integrin^+^/CD9^+^ cells gated were clustered using the X-shift algorithm (K=50 was auto-selected by the switch-point finding algorithm) yielding 61 clusters. Up to 10,000 cells were randomly selected from each X-shift cluster, each cell was connected to 50 nearest neighbors in the phenotypic space and the graph layout was generated using the ForceAtlas2 algorithm (n= 3 per condition). Identified clusters were visualized using single-cell force-directed layout. The expression levels of the myogenic transcription factor Pax7 (upper panels) and the surface marker CD47 (lower panel) were overlaid and are shown in the panel composite. **(E)** The bar graph shows the change in the proportion of each MuSC subset in aged mice treated with THBS1 blockade, relative to IgG control (n= 3 mice per condition). One-tailed paired t-test was used to determine the difference in the proportion of CD47^lo^ and CD47^hi^ MuSCs between IgG control and *α*-THBS1 treatment in aged samples. **(F)** Bar graph shows the fraction of Pax7^+^ cells in the Live/Lineage^-^/*α*_7_integrin^+^/CD9^+^ cell population from young control mice and aged control and treated mice (n= 3 mice per condition). One-tailed unpaired t-test was used to determine the difference in the proportion of Pax7^+^ cells between young and aged IgG control. One-tailed paired t-test was used to determine the difference in the proportion of Pax7^+^ cells between IgG control and *α*-THBS1 treatment in aged samples. **(G)** Experimental scheme. Young and aged mice were pretreated (three times, at two-days intervals) by intramuscular injection in the TA and GA muscles with a blocking antibody to THBS1 or IgG control (contralateral leg) prior to notexin injury and hindlimb muscle tissue was collected for CyTOF analysis 3- or 6-days post injury. **(H)** The scatter plot shows the proportion of MuSCs within the myogenic compartment for each condition, at day 3 post injury (mean ± SEM, n=6, 2 independent experiments). Two-way ANOVA with Sidak’s correction for multiple comparisons was used to determine the difference in the abundance of MuSCs between IgG control and *α*-THBS1 treatment in young or aged samples, and between young and aged IgG treated samples at day 3 post injury. **(I)** Model. Skeletal muscle injury leads to MuSC activation. In young muscle during regeneration, progenitor cells participate in a negative feedback loop whereby they produce THBS1 to limit MuSC proliferation, and promote MuSC return to quiescence, therefore preventing MuSC exhaustion. In aged muscle accumulation of a dysfunctional CD47^hi^ MuSC subset, which precociously secretes THBS1, creates a dysregulated microenvironment that inhibits the proliferation and function of the CD47^lo^ MuSC subset, impairing muscle regeneration. *, *** and **** represent statistical significance at p*≤*0.05, p*≤*0.001 and p*≤*0.0001 respectively.

To determine whether thrombospondin-1 blockade *in vivo* could ameliorate muscle structure and function, we performed histological analysis of skeletal muscle from *Pax7^CreERT2^;Rosa26-LSL-Luc* mice, as well as grip strength 14 days after the end of treatment. Thrombospondin-1 blockade led to a significant increase in Pax7^+^ cells, muscle cross-sectional area and minimum Feret’s diameter compared to IgG treated muscles (Fig. S5 A-E). Moreover, mice treated with the anti-thrombospondin-1 antibody exhibited increased grip strength compared to the IgG treated controls (Fig. S5F).

To investigate the magnitude of MuSC activation and proliferation in young and aged mice upon thrombospondin-1 blockade *in vivo*, we treated mice with the same antibody regimen used above in the TA and GA muscles and employed CyTOF analysis of skeletal muscle to measure IdU incorporation in the previously defined activated MuSC subset(Porpiglia et al., 2017) (CD44^+^/CD98^+^, Fig. 5B). Strikingly, aged mice that received thrombospondin-1 blockade exhibited a two-fold increase in the proportion of activated stem cells and a significant increase in the proportion of proliferating stem cells that incorporated IdU, as shown by color overlay (Fig. 5B, biaxial plots), suggesting that *in vivo* thrombospondin-1 blockade boosts the proliferative capacity of aged MuSCs in resting muscle.

To determine whether activated aged MuSCs returned to quiescence following *in vivo* thrombospondin-1 blockade, hindlimb muscles from aged mice treated with the anti-thrombospondin-1 regimen were collected 6 days after the end of treatment (Fig. 5C). High-resolution CyTOF analysis of the entire myogenic compartment, using X-shift clustering paired with single-cell force directed layout visualization, revealed that aged mice treated with control IgG exhibited a significant decrease in the proportion of Pax7^hi^ stem cells in their TA and GA muscles compared to young mice (Fig. 5D, F). Strikingly, thrombospondin-1 blockade was sufficient to significantly increase the number of Pax7^hi^ stem cells to a level more similar to that seen in young mice (Fig. 5D, F). Moreover, the Pax7^hi^ stem cells in treated aged mice expressed lower levels of CD47 compared to control, suggesting that thrombospondin-1 blockade facilitated the expansion of CD47^lo^ MuSCs in aged mice (Fig. 5D, E). These results show that thrombospondin-1 blockade leads to a remarkable MuSC activation in aged muscle in the absence of injury.

### Thrombospondin-1 blockade *in vivo* enhances the regenerative response of aged muscle

To determine whether thrombospondin-1 blockade could enhance the regenerative response of aged MuSCs *in vivo*, TA and GA muscles of young and aged mice were treated with the anti-thrombospondin-1 regimen described above and then acutely injured by notexin injection. Hindlimb muscles were harvested 3- and 6-days post injury and analyzed by CyTOF to quantify MuSC expansion at day 3 and Pax7 expression at day 6 (Fig. 5G). We found that at day 3 post injury, when MuSC expansion is normally at its peak(Blau et al., 2015; Porpiglia et al., 2017), control IgG-treated aged MuSCs, defined as Lin^-^/*α*_7_integrin^+^/CD9^int^, comprised a smaller fraction of the total myogenic compartment than control IgG treated young MuSCs (Fig. 5H).

Strikingly, *in vivo* thrombospondin-1 blockade led to a significant increase in the proportion of aged MuSCs at day 3 post injury, compared to control (Fig. 5H), as well as an increase in the proportion of aged activated (CD98^+^/CD44^+^) and proliferating (IdU^+^) MuSCs (Fig. S5G), suggesting improved regenerative capacity. Moreover, thrombospondin-1 blockade was sufficient to increase the number of Pax7^hi^ cells in the aged myogenic compartment at day 6 post injury (Fig. S5H), suggesting that the expanded aged MuSC population was able to self-renew. Taken together our data indicate that transient modulation of thrombospondin-1 signaling *in vitro* and *in vivo* by a thrombospondin-1 blocking antibody treatment represents a promising therapy to restore the regenerative potential of aged MuSCs.

## DISCUSSION

The loss of muscle mass and strength with aging is a major predictor of poor health outcome, with a cost of billions of healthcare dollars(Goates et al., 2019; Landi et al., 2012; Rockwood and Mitnitski, 2007; Rolland et al., 2008). Sarcopenia, now recognized as a disease by the World Health Organization(Cao and Morley, 2016), is due in part to the accumulation of dysfunctional stem cells that have lost the ability to proliferate and regenerate the tissue(Bernet et al., 2014; Cosgrove et al., 2014; Price et al., 2014; Tierney et al., 2014). Previous studies from our group and others have identified cell intrinsic and extrinsic signaling pathways that are dysregulated in aged MuSCs, leading to defects in quiescence, self-renewal and proliferation(Bernet et al., 2014; Chakkalakal et al., 2012; Cosgrove et al., 2014; Lukjanenko et al., 2016, 2019; Price et al., 2014; Rozo et al., 2016; Tierney et al., 2014; Vinel et al., 2018). However, a major barrier to elucidating the molecular mechanisms responsible for the age-associated decline in MuSC regenerative capacity has been the heterogeneity of the MuSC population, underscoring the need for single-cell studies. To resolve the molecular and functional heterogeneity of the MuSC population and distinguish dysfunctional subsets that can be purified and characterized during aging, we capitalized on mass cytometry(Bjornson et al., 2013; Spitzer and Nolan, 2016). Here we uncover a new role for the surface marker CD47 as a regulator of skeletal muscle stem cell function in aging. CD47 is a transmembrane protein that belongs to the immunoglobulin superfamily and has been extensively studied in the context of hematopoiesis, where it functions as both a receptor and a ligand(Lindberg et al., 1993, 1994). It is a ligand for SIRP*α*, through which CD47 prevents phagocytosis(Brown and Frazier, 2001), and a receptor for the extracellular matrix protein thrombospondin-1 (Gao et al., 1996; Liu et al., 2001). However, the function of CD47 in stem cell fate in non-hematopoietic tissues has been understudied. Our studies highlight a new role for CD47 levels in determining muscle stem cell fate and function: self-renewal or commitment.

While CD47 is expressed on all cells, its expression levels are transiently altered in different contexts, modulating its function(Jaiswal et al., 2009; Khandelwal et al., 2007; Oldenborg et al., 2000; Reinhold et al., 1995; Van et al., 2012). Previous studies have shown that CD47 expression is increased on the surface of hematopoietic stem cells (HSCs) upon damage-induced mobilization and that this increase protects HSCs in the periphery from being cleared by phagocytes during immune surveillance, a process that is hijacked by cancer cells(Jaiswal et al., 2009; Majeti et al., 2009). Elevated expression of CD47 on CD4 T cells has been shown to define functional long-lived memory T cell precursors(Van et al., 2012). Finally, decreased CD47 levels mediate age-related clearance of red blood cells, providing evidence for changes in CD47 expression with age(Khandelwal et al., 2007). However, a role for CD47 differential expression in muscle stem cells within the context of regeneration and aging has not been previously reported. Here we show that in contrast to aged red blood cells with low CD47 levels, aged MuSCs have elevated CD47 levels, which likely prevent their clearance, and that this dysfunctional MuSC subset, characterized by poor regenerative capacity, accumulates and becomes predominant with age.

We sought to establish the mechanism by which the dysfunctional CD47^hi^ subset arises with aging. We identified an increased abundance of CD47 mRNA isoforms with long 3’UTR in aged MuSCs. We showed that this isoform switch results from alternative polyadenylation, which generates CD47 mRNA isoforms that dictate different protein localization and function. Specifically, we found that in young muscle, expression of the long CD47 mRNA isoform is elevated at the stem to progenitor cell transition, leading to a switch from an intracellular to a cell surface form of CD47 protein. In aged muscle, in contrast to young, the long CD47 mRNA isoform is increased prematurely in MuSCs, leading to the accumulation of a dysfunctional CD47^hi^ MuSC subset. Importantly, we identify U1 snRNA as a key factor responsible for the premature switch to the long CD47 mRNA isoform in aged MuSCs. Strikingly, we found that engraftment of aged MuSCs following transplantation *in vivo* was markedly increased by preventing the binding of U1 snRNA to CD47 3’ UTR using antisense morpholino oligonucleotides, which restored the balance of CD47 mRNA isoforms toward the short isoform, as seen in young MuSCs (Fig. 3G). Previously, alternative polyadenylation was shown to play a role in modulating MuSC function in different muscles, by generating isoforms of Pax3 with different susceptibility to miRNA binding, leading to translational repression(de Morree et al., 2019). Our findings underscore the importance of another mechanism, protein localization, by which post-transcriptional control regulates muscle stem cell fate.

To understand the role of CD47 in the regulation of MuSC fate and function we investigated its interaction with thrombospondin-1, a negative regulator of cell proliferation, as this pathway has previously been implicated in self-renewal and reprogramming(Kaur et al., 2013; Soto-Pantoja et al., 2015). Moreover, thrombospondin-1 upregulation during the course of aging has been previously reported in several tissues, including kidney, heart and skeletal muscle(Hoier et al., 2013a, 2013b; Isenberg and Roberts, 2020; Kivelä et al., 2008). We found that in aged muscle tissue, in the context of injury, CD47^hi^ MuSCs secrete high levels of thrombospondin-1, which inhibits the expansion of the residual functional CD47^lo^ MuSC subset, in a paracrine loop leading to global impairment of aged MuSC proliferation and regeneration. Using live single-cell confocal imaging of sorted MuSC subsets expressing a fluorescent cAMP sensor, we showed that thrombospondin-1-mediated suppression of cell proliferation occurs through inhibition of cAMP production. Importantly, we demonstrate that transient thrombospondin-1 blockade in two different *in vivo* contexts, in acute muscle injury as well as in resting muscle in the absence of injury, is sufficient to promote the activation and self-renewal of the aged CD47^lo^ MuSC subset and markedly enhance aged muscle regenerative response. Intriguingly, in young muscles undergoing hypertrophy in response to mechanical stress, thrombospondin-1/CD47 signaling induced by an exogenous thrombospondin-1 peptide was found to promote MuSC proliferation via a mechanism that requires calcitonin receptor downregulation and is independent of cAMP(Kaneshige et al., 2022). Our findings underscore the complexity of CD47 signaling and reveal an age-dependent alteration in the role of CD47 in MuSC function.

Our work suggests that cancer immunotherapy may impact cancer cachexia. CD47 upregulation has been identified as a means by which cancer cells escape immune clearance, by engaging SIRP*α* on immune cells(Jaiswal et al., 2009; Majeti et al., 2009). This interaction is currently being targeted systemically for cancer immunotherapy(Feng et al., 2019; Majeti et al., 2009). Further investigation is warranted to determine if immunotherapies that interfere with thrombospondin-1/CD47 signaling in peripheral tissues such as muscle, could lead to MuSC activation and increased stem cell proliferation and counteract the muscle wasting that accompanies cancer.

In summary, the work presented here reveals a novel aging-associated mechanism underlying MuSC dysfunction and a means of isolating the dysfunctional aged MuSC subset. The purification of defective aged MuSCs enabled unprecedented mechanistic insights and allowed us to design studies to overcome aged MuSC dysfunction and ameliorate muscle regeneration in aging *in vivo* by two complementary approaches: (i) decreasing CD47 expression by altering polyadenylation site choice via delivery of antisense morpholino oligonucleotides or (ii) antibody blockade of thrombospondin-1/CD47 signaling. These findings have therapeutic potential and are of broad significance to the fields of aging and regenerative medicine.

## ACKNOWLEDGEMENTS

We thank Kassie Koleckar for technical assistance, Drs. David Burns, Glenn Markov, Megan Mayerle and Gaspard Pardon for valuable discussions, the Stanford Shared FACS Facility for technical support, Drs. Anthony Jose and Nori Ueno from Thermo Fisher Scientific for support with the PrimeFlow RNA assay, and Dr. Miro Koulnis for help with illustrations. We thank Dr. Garry P. Nolan for his expertise and critical insight into single-cell analysis and for sharing resources. This study was supported by BD Biosciences Stem Cell grant (E.P.); NIH grants NS089533 and AG020961, California Institute for Regenerative Medicine grant RB5-07469 and the Baxter Foundation (H.M.B.).

## AUTHOR CONTRIBUTIONS

E.P. and H.M.B. conceived the study. E.P. designed and performed experiments, analyzed and interpreted data and wrote the manuscript. H.M.B and W.J.F. designed experiments, analyzed and interpreted data and wrote the manuscript. T.M. and A.D.M. designed and performed experiments, analyzed and interpreted data. P.K. performed experiments and analyzed data. C.A.H. developed bioinformatic tools for image processing and performed segmentation analysis of histological section images. S.B. performed analysis of histological section images. V.D.G., and P.K.J. analyzed and interpreted data. K.H. contributed to discussions of data and the cAMP assay in living cells. L.F. performed bioinformatic analysis of published RNA-seq datasets. A.T. provided technical support with antibody conjugation and CyTOF data acquisition.

## METHODS

### Mouse lines

All animal protocols were approved by the Stanford University Administrative Panel on Laboratory Animal Care (APLAC) (protocol #10509 and #29398) and experiments were performed in compliance with the institutional guidelines of Stanford University. C57BL/6 young adult mice were purchased from Jackson Laboratories (Bar Harbor, ME) and used at 2-4 months of age (median age 2 months). Aged C57BL/6 mice (> 22 months) were purchased from the US National Institute on Aging (NIA). Mice ubiquitously expressing a green fluorescent protein (GFP) transgene and mice ubiquitously expressing a firefly luciferase (Fluc) transgene driven by the ACTB promoter (L2G85 strain) were obtained and genotyped as described previously(Sacco et al., 2008). Double transgenic GFP/Luciferase mice were generated by breeding the above strains and were confirmed by appropriate PCR-based strategies to validate the genotype. CD47^-/-^ mice(Lindberg et al., 1996) (Stock # 003173) were obtained from Jackson Laboratory and genotyped according to the recommended PCR-based protocol. Cells from C57BL/6 mice and CD47^-/-^ mice were used for cell proliferation and protein expression experiments. Cells from GFP/Luciferase mice were isolated for transplantation experiments from young or aged mice. Double-transgenic *Pax7^CreERT2^;Rosa26-LSL-Luc* were generated by crossing *Pax7^CreERT2^* mice obtained from Jackson Laboratory (Stock # 017763) and *Rosa26-LSL-Luc* obtained from Jackson Laboratory (Stock # 005125). We validated these genotypes by appropriate PCR-based strategies. The number of animals for each data set and all relevant details regarding the sample size are reported with each experiment.

### Cell isolation and dissociation of muscle tissue

*Tibialis Anterior* (TA) and *Gastrocnemius* (GA) muscles were dissected and subjected to mechanical dissociation using the gentleMACS^TM^ dissociator (Miltenyi Biotech), followed by collagenase (0.25%) and dispase (0.04 U/ml; Roche, Indianapolis, IN) digestion at 37°C for 90 minutes. The resulting cell suspension was passed through a standard syringe needle and subsequently through a 70-μm nylon filter (BD Biosciences, San Jose, CA). No established cell lines were used in this study.

### Isolation of muscle stem cells and their subsets

Muscle tissue was isolated and digested to a single-cell suspension as described above. Cells were first incubated with a purified antibody reactive to murine CD47 (Biolegend, San Diego, CA) for 30 minutes at 4°C, then washed and incubated with an anti-Rat IgG2a secondary antibody conjugated to Percp-eFluor710 (eBioscience, San Diego, CA). Cells were then incubated with biotinylated antibodies reactive to murine CD45, CD11b, CD31, and Sca1 (BD Biosciences, San Jose, CA) for 30 minutes at 4°C and washed. Cells were then incubated with streptavidin magnetic beads (Miltenyi Biotech, Auburn, CA), streptavidin-APC-Cy7 (Invitrogen, Carlsbad, CA), *α*7 integrin-PE antibody (AbLab, Vancouver, Canada), CD34-eFluor660 antibody (eBioscience, San Diego, CA). After magnetic depletion of the biotin-positive cells, the lineage-negative cells were stained with DAPI and sorted on a FACSAria cell sorter or a Sony SH800 cell sorter in purity mode, using FACS Diva software (BD Biosciences, San Jose, CA) or SH800 software (Sony). To isolate MuSCs, we first gated on viable cells (DAPI negative), then on cells negative for the lineage markers (CD45, CD31, CD11b, and Sca1), finally on cells positive for both CD34 and *α*7 integrin, which represent the muscle stem cell fraction. To simultaneously isolate CD47^lo^ and CD47^hi^ subsets we first gated on viable cells (DAPI negative), then on cells negative for the lineage markers (CD45, CD31, CD11b, and Sca1) and positive for *α*7 integrin and CD34. Finally, we sorted CD47^lo^ and CD47^hi^ subsets based on their increasing CD47 expression. Flow cytometry scatter plots were generated using FlowJo v8.7 (Treestar, Ashland, OR).

### Biomimetic Hydrogel Fabrication

We fabricated polyethylene glycol (PEG) hydrogels from PEG precursors, synthesized as described previously(Gilbert et al., 2010). We produced hydrogels by using the published formulation to achieve 12-kPa (Young’s modulus) stiffness hydrogels in 1-mm thickness, which is the optimal condition for culturing MuSCs and maintaining stem-cell fate in culture(Gilbert et al., 2010).

### Transplantation of stem cells and their subsets

Sorted cell populations were transplanted immediately following FACS isolation into the TA muscle of recipient mice as previously described(Sacco et al., 2008), or 16 hours post treatment with either antisense morpholino oligonucleotides, as described below. Cells from GFP/Luciferase mice were transplanted into gender-matched, hindlimb-irradiated NOD/SCID mice (Jackson Laboratories, Bar Harbor, ME). NOD/SCID mice (2-4 months of age, median 2 months) were anesthetized with ketamine (2.4 mg per mouse) and xylazine (240 μg per mouse) by intraperitoneal injection and irradiated by a single dose of 18 Gy administered to the hindlimbs, with the rest of the body shielded in a lead jig. Transplantation was performed within three days post-irradiation. Freshly isolated cells were counted by hemocytometer and resuspended at desired cell concentrations in PBS with 2.5% goat serum and 1 mM EDTA. Cells were transplanted by intramuscular injection into the TA muscle in a 10 μl volume. To evaluate stem cell repopulation, we collected and digested primary recipient muscles four weeks after transplant and stained them with antibodies to lineage markers (CD45, CD11b, CD31 and Sca1), surface markers *α*7 integrin and CD34, as well as with DAPI, to determine viability. To identify donor cells, we gated for cells negative for DAPI and the lineage markers and positive for *α*7 integrin and CD34 and the donor marker GFP.

### *In vivo* IdU labeling

Mice were weighed, anesthetized with isofluorane and injected intraperitoneally with IdU eight hours prior to sacrificing the animal (150 mg/kg body weight per injection).

### Antisense Morpholino Oligonucleotides treatment

Freshly sorted MuSCs were plated in Ham’s F10, 10% Horse Serum and 1% Penicillin/Streptomycin at 37C in 24 well plates and transfected with antisense morpholino oligonucleotides (AMO) at a concentration of 15μM with Endoporter (Gene Tools, LLC), according to the manufacturer’s instructions. For RNA preparation, media was changed 16 hours post transfection and samples were collected at 24 hours post transfection. For flow cytometric analysis of CD47 expression, media was changed 16 hours post transfection and samples were collected at 32 hours post transfection. For transplantation assays cells were collected 16 hours post transfection and transplanted as described above.

We used the following AMO sequences:

Control AMO: ggttacaatctaagatcaaacgacg; CD47_PAS1 AMO: cacagcacatcatattttttttatt; CD47_U1 AMO: agtacagctgctttgaatataaact

### Histology

We collected and prepared TA muscle tissue for histology as previously described(Sacco et al., 2008). We fixed transverse sections from muscles using 4% PFA, blocked and permeabilized using PBS/1% BSA/0.1% Triton X-100 and incubated with anti-Laminin (Abcam, cat # ab11575, 1:500), anti-PAX7 (Santa Cruz Biotechnology, cat# sc-81648, 1:50) and then with AlexaFluor secondary Antibodies (Jackson ImmunoResearch Laboratories, 1:200) We counterstained nuclei with DAPI (Invitrogen). Images were acquired using KEYENCE BZ-X700 all in one fluorescence microscope with 20x/0.75 N.A. objective. Image processing of the skeletal muscle immunofluorescent histological sections, for semi-automated fiber detection and analysis of fiber cross-sectional area and minimum feret’s diameter, was performed using the SMASH plugin for MATLAB(Smith and Barton, 2014). Pax7^+^ cells were quantified manually using ImageJ. Data analyses were blinded.

### Grip strength measurements

Mice were put on a BioSeb grip tester and their grip strength was measured for one hindlimb at the time(Serrano and Muñoz-Cánoves, 2021). Mice were measured three-six times and the average was used to calculate grip strength. Values were normalized to mouse weight or to the IgG control leg.

### Bioluminescence imaging

Bioluminescence imaging (BLI) was performed using a Xenogen-100 system, as previously described(Sacco et al., 2008), at 3-4 weeks post-transplant or before and after *in vivo* anti-thrombospondin-1 treatment for *Pax7^CreERT2^;Rosa26-LSL-Luc*. The system is comprised of a light-tight imaging chamber, a charge-coupled device (CCD) camera with a cryogenic refrigeration unit and the appropriate computer system (Living-Image Software, Caliper LifeSciences). Briefly, the animals were anesthetized under isofluorane and injected intraperitoneally with a 100μl volume of luciferin diluted in PBS (0.1mmol/Kg body weight, Caliper LifeSciences). For transplantation, immediately after injection, images were acquired each minute for a total of 15 min and data were stored for subsequent analysis, using Living Image software (Caliper LifeSciences). Bioluminescence images acquired at 12 min post-luciferin injection were used for analysis. Bioluminescence signal was calculated by drawing a consistent region-of-interest (ROI) over each hindlimb and quantifying the resulting signal. A bioluminescence signal value of 80,000 photons/s represented our detection threshold, as this level was previously determined to represent the presence of one or more GFP-positive muscle fiber in transplanted tissue. For measuring endogenous muscle stem cell expansion, immediately after injection, images were acquired at 1, 5, 30, 60, 120 seconds and data were stored for subsequent analysis, using Living Image software (Caliper LifeSciences). Bioluminescence images acquired at 30 seconds post-luciferin injection were used for analysis as described above.

### Muscle injury

Mice (8-10 weeks) were acutely injured by a single 10 μl intramuscular injection of notexin (10 μg/ml; Latoxan, France) into the TA muscle and two injections in the GA muscle at the indicated time points.

### Endogenous muscle stem cell lineage tracing

*Pax7^CreERT2^; Rosa26-LSL-Luc* mice were treated with five consecutive daily intraperitoneal injections at a tamoxifen dose of 75mg/kg body weight to activate luciferase expression following *Pax7-*dependent Cre-mediated recombination. Ten days after the last tamoxifen injection, mice were subjected to repeated anti-thrombospondin-1 injections (3 injections, at two days intervals) and imaged by BLI.

### Surface and intracellular flow cytometry staining for CD47

Muscle tissue was isolated and digested to a single-cell suspension as described above. Cells were incubated with biotinylated antibodies reactive to CD45, CD11b, CD31, and Sca1 (BD Biosciences, San Jose, CA) for 30 minutes at 4°C and washed. Cells were then incubated with streptavidin-APC-Cy7 (Invitrogen, Carlsbad, CA), *α*7 integrin-PE antibody (AbLab, Vancouver, Canada), CD9-APC antibody (eBioscience, San Diego, CA) and CD47-BV605 antibody (BD Biosciences, San Jose, CA) for 30 minutes on ice and washed. Cells were fixed in 1.6% paraformaldehyde, permeabilized with BD permwash buffer I (cat# 557885, BD Biosciences, San Jose, CA), washed and stained intracellularly in for CD47-PECy7 (Biolegend, San Diego, CA) in BD permwash buffer I. Finally, cells were washed twice with staining buffer and analyzed on a BD-LSRII flow cytometer using FACS Diva Software (BD Biosciences, San Jose, CA).

### PrimeFlow RNA assay

RNA in situ hybridization probes were custom-designed by Thermo Fisher Scientific to differentiate between the short and long 3’UTR isoforms of CD47 mRNA. About 8000 fluorophores labeled each target RNA. TA and GA muscles were isolated from young (2 months) and aged (24 months) mice and digested to single cell suspensions that were stained using antibodies against lineage markers (CD45, CD11b, CD31, Sca1)-APC-Cy7, *α*7 integrin-PE, CD9-APC and CD47-BV605. Samples were then processed for the PrimeFlow RNA Assay (cat # 88-18005-204, Thermo Fisher Scientific), following the manufacturer’s instructions. Cells were fixed, permeabilized, and stained for intracellular CD47 using CD47-PE-Cy7 antibody. Finally, RNA in situ hybridization was performed using custom-designed probes targeting the total (Assay ID: VB1-3030381) or long (Assay ID: VB6-6001206) 3’UTR isoform of CD47 mRNA. The total and long 3’UTR isoforms of CD47 mRNA were labeled with AF647 and AF750, respectively.

### MuSC culture and treatment

Following isolation, we resuspended MuSCs in myogenic cell culture medium containing DMEM/F10 (50:50), 20% FBS, 2.5 ng·ml^−1^ fibroblast growth factor-2 (FGF-2, also known as *β*FGF) and 1% penicillin–streptomycin. We seeded MuSC suspensions at a density of 500 cells per cm^2^ surface area. We maintained cell cultures at 37°C in 5% CO_2_ and changed medium every other day. For thrombospondin-1 blockade studies, we added the thrombospondin-1 blocking antibody (10μg/ml) (A6.1, Thermo Fisher Scientific) to MuSCs cultured on biomimetic hydrogels 24 hours post isolation and cultured the cells for a total of 6 days, changing the media every two days. Cell number was determined by manual cell counting. For thrombospondin-1 treatment studies (dose response curve), we added recombinant thrombospondin-1 (R&D Systems) at the indicated concentrations 24 hours post isolation and cultured the cells for a total of 6 days, changing the media every two days. Cell number was determined by manual cell counting.

### *In vitro* IdU incorporation

MuSCs cultured on biomimetic hydrogel and treated with thrombospondin-1 or with the thrombospondin-1 blocking antibody (A6.1, Thermo Fisher Scientific) for 6 days were pulsed with IdU at a concentration of 10μm for 1 hour and then harvested and processed for CyTOF staining as described above.

### Live Cell cAMP assay

CD47^lo^ and CD47^hi^ MuSC subsets were sorted and seeded at 3000 cells/well in a 96-well cell imaging plate (Eppendorf, 0030741013) and transduced the following day with the cADDis BacMam sensor (Molecular Montana, D0200G) according to manufacturer’s recommendation(Tewson et al., 2016). Briefly, cell were infected with 10μl of BacMam sensor stock in a total of 150ul of media containing 2mM Sodium Butyrate (Molecular Montana) for 30 min at room temperature followed by 6 hours in the 37°C tissue culture incubator. BacMam was removed and replaced with media containing 1mM Sodium Butyrate for 16-24 hours. Prior to imaging, cells were incubated in PBS for 20 min at room temperature. Images were acquired on a Marianas spinning disk confocal (SDC) microscope (Intelligent Imaging Innovations) (40x, epi-fluorescence) every 10 seconds for 5 min, with thrombospondin-1 (5μg/ml) added after 30 seconds. Red fluorescence was used to determine a mask and background subtracted green and red fluorescent intensity over time was determined using Slidebook (Intelligent Imaging Innovations).

### cAMP competitive ELISA

Sorted MuSCs were cultured overnight in 96 well plates and cAMP measurements were performed on cell lysates using the Cyclic AMP XP^®^ Assay Kit (cat Cyclic AMP XP^®^ Assay Kit (cat # 4339, Cell Signaling Technology) according to the manufacturer’s instructions.

### Antibody conjugation with metal isotopes

Purified antibodies were conjugated to the indicated metals for mass cytometry analysis using the MaxPAR antibody conjugation kit (Fluidigm, South San Francisco, CA) according to the manufacturer’s instruction. Following labeling, antibodies were diluted in Candor PBS Antibody Stabilization solution (Candor Bioscience GmbH, Wangen, Germany) to 0.2 mg/ml and stored long-term at 4°C. Each antibody clone and lot was titrated to optimal staining concentrations using murine myoblasts, as well as murine muscle and spleen cell suspensions.

### Mass cytometry staining

Dead cells were stained using cisplatin (WR International, Radnor, PA, Cat# 89150-634) as previously described(Fienberg et al., 2012). In particular, cells were resuspended in 1ml serum-free DMEM at 2×10^6^ cells/ml and cisplatin was added at a final concentration of 25 μM for 1 minute at room temperature. The reaction was quenched with 3ml of DMEM/10% FBS. Samples were centrifuged at 1400 rpm for 10 minute and cell pellets were resuspended in cell staining media (CSM: PBS with 0.5% bovine serum albumin (BSA) and 0.02% sodium azide) and fixed with 1.6% paraformaldehyde (PFA) for 10 minutes on ice. Cells were washed with CSM and stained with antibodies against surface markers included in the mass cytometry panel for 1 hour at room temperature. Cells were washed twice with CSM and permeabilized with methanol for 10 minutes on ice. Cells were washed twice with CSM and stained with antibodies against intracellular markers included in the mass cytometry panel for 1 hour at room temperature. Cells were washed twice with CSM and stained with 1 ml of 191/193Ir DNA intercalator (Fluidigm, South San Francisco, CA) diluted (1:5000) in PBS with 1.6% PFA for 20 minutes at room temperature.

### Mass cytometry measurement

Cells were acquired on the CyTOF 2 mass cytometer (Fluidigm, South San Francisco, CA) at an event rate of approximately 500 cells per second as previously described. The instrument was run in high-resolution mode (Mass resolution ∼700) with internally calibrated dual-count detection. Noise reduction and cell extraction parameters were: cell length 10-65, lower convolution threshold 10. Samples were normalized using beta beads.

### X-Shift analysis and graphic display of single-cell mass cytometry data

Events were clustered based on a combination of surface markers and myogenic transcription factors using the X-Shift algorithm and clusters were validated. In order to visualize the spatial relationships between the cell types within these X-shift clusters, 2,000 or 10,000 randomly sampled cells from each cluster were subjected to a force-directed layout. All conditions from a single experiment were processed simultaneously so that the resulting map would capture all populations present in the entire dataset.

### Bulk RNA-Seq

α7-integrin^+^/CD34^+^ MuSCs were isolated as described above. RNA was isolated using Qiagen RNAEasy Micro kit from 5,000-10,000 cells and cDNA generated and amplified using NuGEN Ovation RNA-Seq System v2 kit. Libraries were constructed from cDNA with the TruSEQ RNA Library Preparation Kit v2 (Illumina) and sequenced to 30–40 × 10^6^ 1 × 75-bp reads per sample on a HiSEQ 2500 from the Stanford Functional Genomics Facility, purchased using NIH S10 Shared Instrument Grant S10OD018220. For analysis, RNA sequences were aligned against the *Mus musculus* mm10 reference genome using the Spliced Transcripts Alignment to a Reference (STAR) software. RNA-Seq by Expectation-Maximization (RSEM) software package was used for annotating reads to transcripts. RNA abundance was quantified into transcripts per million (TPM) values as well as total read counts. A counts matrix containing the number of read counts for each gene and each sample was obtained. This matrix was analyzed by DESeq to calculate statistical analysis of significance of genes between samples.

### Single-Cell RNA-seq

Following sorting, cells were counted and diluted to 500 cells per microliter and subsequently loaded, captured and stained with viability dyes (LIVE/DEAD cell viability assay; Molecular Probes, Life Technologies) on a small-sized microfluidc RNA-seq chip (Fluidigm) using the C1 system. Captured cells were imaged using phase-contrast and fluorescence microscopy to determine live/dead status. Lysis, reverse transcription and cDNA preamplification was performed using the SMARTer Ultra Low RNA Kit for the Fluidigm C1 System (Clontech) according to Fluidigm’s manual for mRNA sequencing on the C1 system. Resulting cDNA was harvested and analyzed on the Fragment analyzerTM automated fragment analyzer by Advanced Analytical. Cells with a concentration higher than 0.05 ng/ul were selected for library preparation. Library preparation was performed using Nextera XT DNA Sample Preparation Kit (Illumina) as described in the Fluidigm manual. Following library preparation, cells were pooled and sequenced on an Illumina NextSeq instrument using 2×75 paired end reads on a NextSeq high output kit (Illumina). Reads were mapped to the *Mus musculus* mm10 reference genome using STAR. RSEM was used for determining transcript abundance (TPM). A counts matrix containing the number of counts for each gene and each sample was obtained. This matrix was analyzed by DESeq to calculate statistical analysis of significance of genes between samples.

### Quantitative RT-PCR

We isolated RNA from muscle stem or progenitor cells using the RNeasy Micro Kit (Qiagen). We reverse-transcribed cDNA from total mRNA from each sample using the High-Capacity cDNA Reverse Transcription Kit (Applied Biosystems). We subjected cDNA to RT-PCR using TaqMan Assays (Applied Biosystems) or SYBR Green PCR master mix (Applied Biosystems) in an ABI 7900HT Real-Time PCR System (Applied Biosystems). We cycled samples at 95 °C for 10 min and then 40 cycles at 95 °C for 15 s and 60 °C for 1 min. To quantify relative transcript levels, we used 2−ΔΔCt to compare treated and untreated samples and expressed the results relative to *Rnu2* or *Gapdh* (Fig. 3B, Fig. S4D). Transcript levels for the long isoform of CD47 were expressed relative to *Cd47 total* levels (Fig. S2H, Fig. 3C).

For SYBR Green qRT-PCR, we used the following primer sequences: *Gapdh_*Fw: aggtcggtgtgaacggatttg; *Gapdh_*Rev: tgtagaccatgtagttgaggtca *CD47total_*Fw: cctgttctggggaaagtttgg*; CD47total_*Rev: aggatggctccaacaaccac

*CD47long_*Fw: gatcggaggcatgacaaagc*; CD47long_*Rev: gcaacaaagagacaagaggcg

*Rnu1*_Fw: cttacctggcaggggagata; *Rnu1*_Rev: atccggagtgcaatggataa

*Rnu2*_Fw ctcggccttttggctaagat; *Rnu2*_Rev tgtcctcggatagaggacgta

TaqMan Assays (Applied Biosystems) were used to quantify *cdkn1a*, *cdkn1b*, *cdkn1c* in samples according to the manufacturer instructions with the TaqMan Universal PCR Master Mix reagent kit (Applied Biosystems). Transcript levels were expressed relative to *Gapdh levels.* Multiplex qPCR enabled target signals (FAM) to be normalized individually by their internal *Gapdh* signals (VIC).

### Statistics and Reproducibility

All animal experiments were randomized. No statistical method was used to predetermine sample size. The investigators were not blinded to allocation during experiments and outcome assessment, except for force measurement assays and histological analyses. Transplantation experiments were performed with at least n=12 mice, two independent experiments. The line indicated in the scatter plots or bar graphs represents the median (Fig. 1G; Fig. 3F), mean ± SEM (Fig. 1C, E; Fig. S1B, C; Fig. 2B, C, F, G; Fig. S2B, G, H; Fig. 3B-E; Fig. S3A-D; Fig. 4A, B, E, F, H, K, L; Fig. S4D-F, H, I; Fig. 5A-B, H; Fig. S5B, D-H) or mean ± SD (Fig. S1D; Fig. 4C, D; Fig. S4A-C; Fig. 5F). ANOVA or multiple t-test analysis were performed for experiments with 3 or more groups, as indicated in the legend. For comparisons between two groups, unpaired or paired Student’s t-test was used. Analyses were conducted using GraphPad Prism 9 software (GraphPad Software). Statistical significance was determined by p*≤*0.05. *, **, *** and **** represent statistical significance at p*≤*0.05, p*≤*0.01, p*≤*0.001 and p*≤*0.0001, respectively.

### Code Availability

The computational code used in the study can be obtained at https://github.com/nolanlab/vortex/releases.

### Data Availability

CyTOF data that support the findings of this study will be deposited in Flowrepository.org RNAseq datasets will be made available upon publication.

### Supplemental information and data

**Fig. S1.**
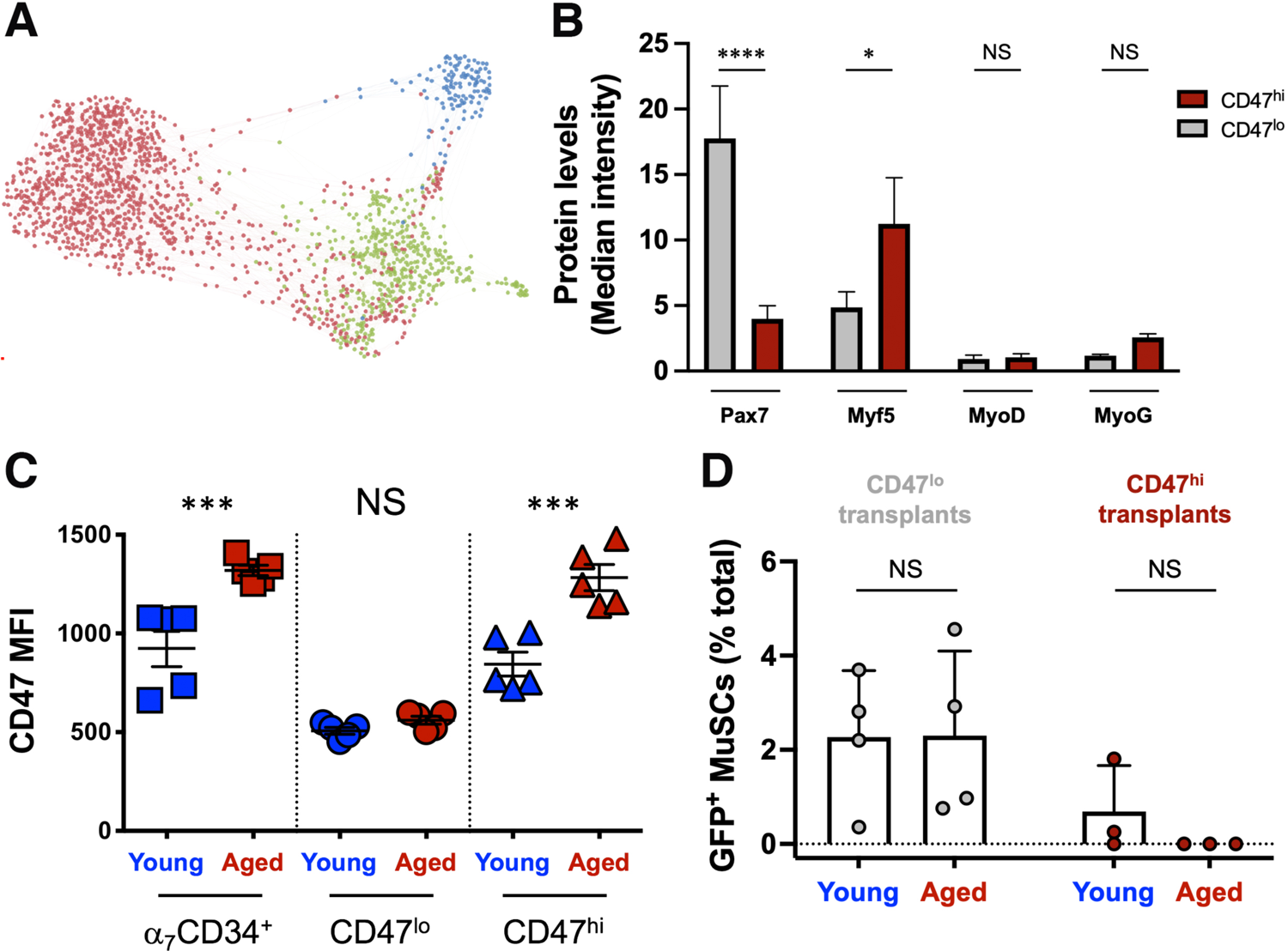
**(A)** CyTOF analysis was performed on TA and GA hindlimb muscle. Gated Live/Lin^-^/*α*_7_integrin^+^/CD34^+^ MuSCs were analyzed with the X-shift algorithm (K=30 was auto-selected by the switch-point finding algorithm) yielding 3 clusters (colour-coded in red, green and blue) that were visualized using single-cell force-directed layout. Up to 2000 cells were randomly selected from each X-shift cluster, each cell was connected to 30 nearest neighbors in the phenotypic space and the graph layout was generated using the ForceAltas2 algorithm as previously described(Porpiglia et al., 2017) (representative experiment, n= 3 mice; 4 independent experiments). **(B)** Quantification of the expression level of myogenic transcription factors Pax7, Myf5, MyoD and Myogenin in CD47^lo^ and CD47^hi^ MuSC subsets, measured by CyTOF analysis as median intensity (mean ± SEM from n=9 mice, 3 independent experiments). Two-way ANOVA analysis with Sidak’s correction for multiple comparisons was used to determine difference in transcription factor expression between the two MuSC subsets. **(C)** Quantification of CD47 protein expression, measured by flow cytometry as median fluorescence intensity (MFI), in young (2 months) and aged (24 months) MuSCs (Lin^-^/ *α*_7_integrin^+^/CD34^+^), CD47^lo^ and CD47^hi^ MuSC subsets (mean ± SEM from n=9 mice, 3 independent experiments). Two-way ANOVA analysis with Sidak’s correction for multiple comparisons was used to determine difference between young and aged subsets. **(D)** Stem cell repopulation analysis in primary recipients performed by flow cytometric detection of donor-derived (GFP^+^) MuSCs as a fraction of the total recipient MuSC (α_7_ integrin ^+^/CD34^+^) population, 4 weeks after transplantation (mean ± SD, n= 4 young and aged CD47^lo^ transplants, n= 3 young and aged CD47^hi^ transplants). Two-way ANOVA analysis with Sidak’s correction for multiple comparisons was used to determine difference between young and aged transplants. *, *** and **** represent statistical significance at p*≤*0.05, p*≤*0.001 and p*≤*0.0001 respectively.

**Fig. S2.**
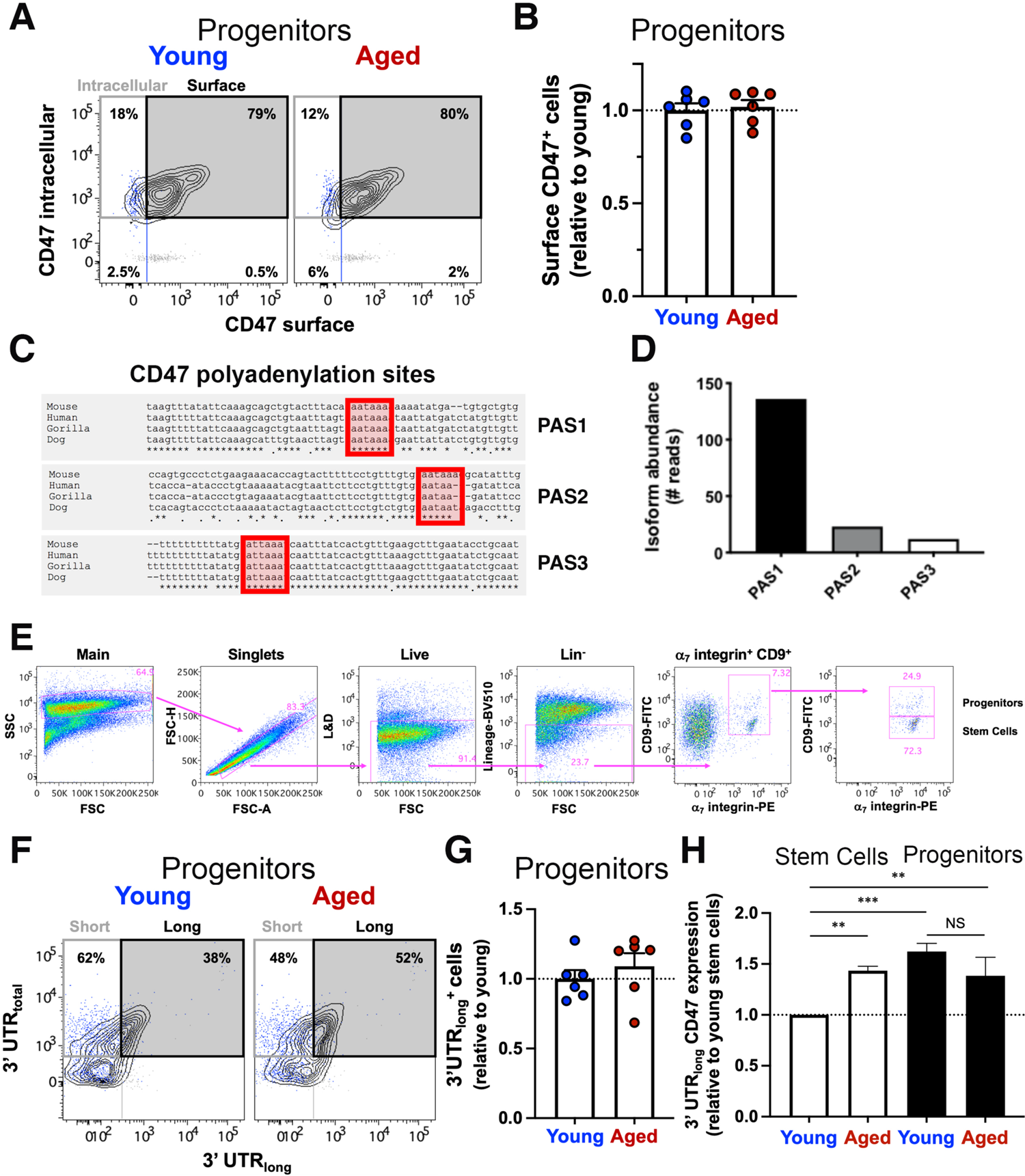
**(A)** TA and GA muscles were isolated from young and aged mice and digested to a single cell suspension that was stained using antibodies against lineage markers (CD45, CD11b, CD31, Sca1)-APCCy7, *α*7 integrin-PE, CD9-APC and CD47-BV605. Cells were then fixed, permeabilized and stained for CD47 intracellularly using a different conjugate CD47-PECy7. A biaxial dot plot of intracellular CD47-PECy7 (y axis) by surface CD47-BV605 (x axis) is shown depicting the distribution of CD47 expression on the surface or inside the cell in progenitor cells from young (left panels) and aged (right panels) mice (representative sample). Overlaid blue dots in the biaxial plot indicate the FMO – surface CD47 control, whereas the overlaid gray dots indicate the FMO – intracellular CD47 control. **(B)** Surface CD47^+^ cells in (A) are quantified. Bar graph indicates the abundance of aged surface CD47^+^ progenitor cells relative to young (mean ± SEM, n=6 mice, 2 independent experiments). Two-way ANOVA analysis was used to determine difference between young and aged subsets. **(C)** The mRNA sequence of CD47 from mouse, human, gorilla and dog were aligned using MAFFT online. 3 highly conserved polyadenylation sites (PAS) were identified (red boxes). **(D)** Quantification of murine CD47 mRNA isoform abundance by analysis of publicly available datasets obtained by 3’ region extraction and deep sequencing. Bar graph shows the abundance of the different CD47 mRNA isoforms. Polyadenylation site 1 (PAS1) = chr16:49911190-49911205; PAS2 = chr16:49,912530-49,912555; PAS3 = chr16: 49,915032-49,915123. **(E)** Gating strategy used to identify stem and progenitor cells on flow cytometry samples described in Fig. 2E-G and Fig. S2F, G. Individual dot plots are shown. **(F)** TA and GA muscles were isolated from young (2 months) and aged (24 months) mice and digested to a single cell suspension that was stained using antibodies against lineage markers (CD45, CD11b, CD31, Sca1)-APCCy7, *α*7 integrin-PE, CD9-APC and CD47-BV605. Cells were then fixed, permeabilized and stained for CD47 intracellularly using a different conjugate CD47-PECy7. Finally, cells were stained according to the PrimeFlow RNA protocol for the different CD47 mRNA isoforms. A biaxial dot plot of CD47 mRNA-3’UTR_total_ (y axis) by CD47 mRNA-3’UTR_long_(x axis) is shown depicting the distribution of cells expressing the CD47 mRNA short isoform (upper left quadrant) or the long isoform (upper right quadrant) as a fraction of total CD47 mRNA (upper left and right quadrants combined) in progenitor cells from young (left panels) and aged (right panels) mice (representative sample). Overlaid blue dots in the biaxial plot indicate the FMO – CD47 mRNA 3’ UTR_long_ control, whereas the overlaid gray dots indicate the FMO – CD47 mRNA 3’ UTR_total_ control. **(G)** Bar graph indicates the relative increase in the abundance of aged progenitor cells expressing the long CD47 mRNA isoform compared to young progenitor cells (mean ± SEM, n=6 mice, 2 independent experiments). 2 tailed paired t-test analysis was used to determine difference between young and aged stem cells. **(H)** Bar graph shows the relative abundance of the long CD47 mRNA isoform, measured by RT-qPCR in young and aged sorted muscle stem and progenitor cells, normalized to young stem cells (mean ± SEM, n=3 for each young and aged population, 2 independent experiments). Two-way ANOVA analysis with Tukey’s correction for multiple comparisons was used to determine difference in the abundance of the long CD47 mRNA isoform across the different conditions in young or aged sorted muscle stem and progenitor cells. ** and *** represent statistical significance at p*≤*0.01 and p*≤*0.001 respectively.

**Fig. S3.**
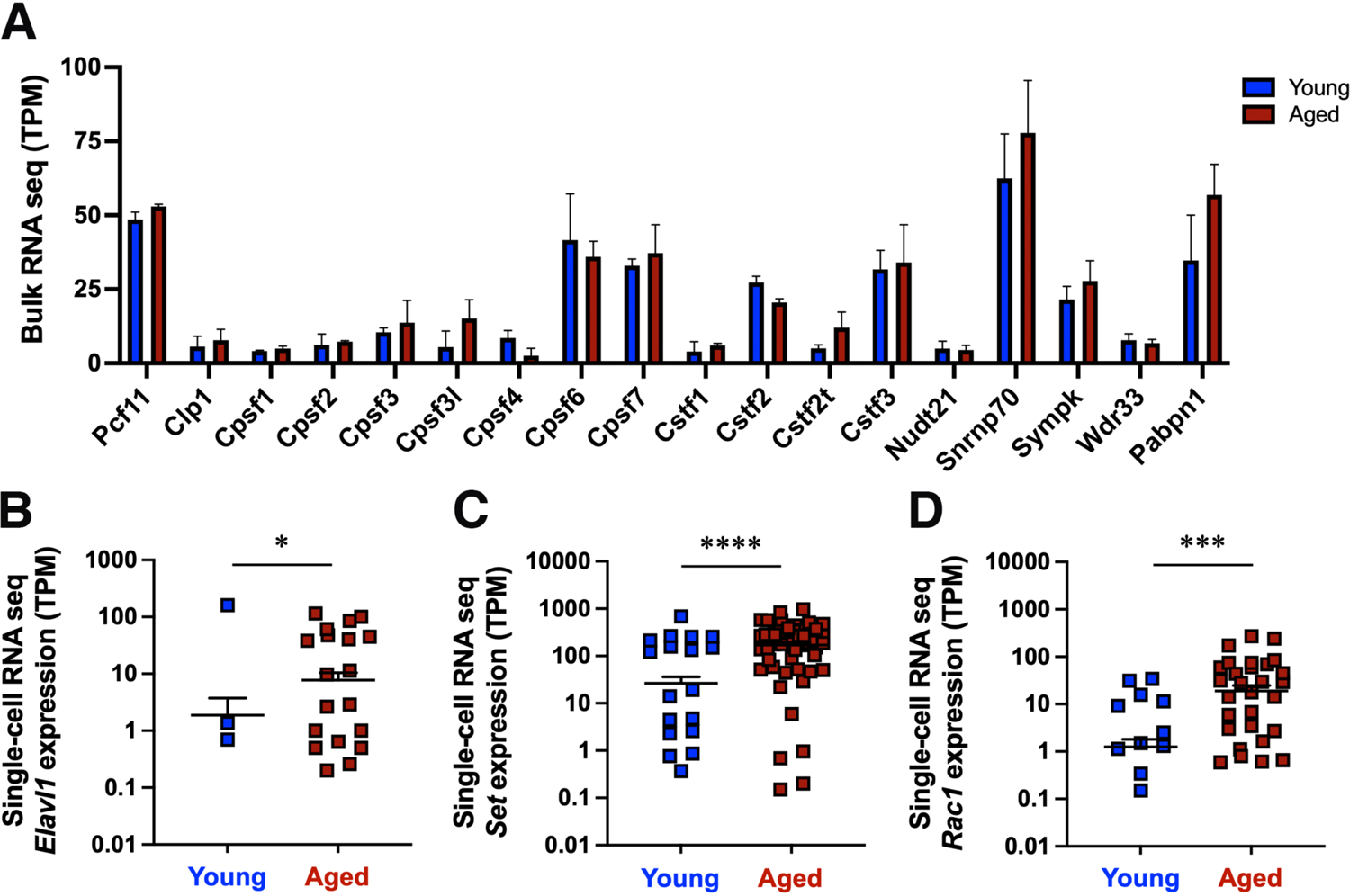
**(A)** Expression of components of the cleavage and polyadenylation factor complex or factors implicated in proximal polyadenylation site readthrough in young and aged MuSCs, measured by bulk RNA seq analysis (mean ± SEM, n=3 young mice, n=3 aged mice). Two-tailed unpaired t-test analysis with Welch’s correction was used to determine the difference in transcript expression between young and aged MuSCs. (**B-D**) Expression of components of the protein complex that traffics the CD47 protein to the cell surface (Elavl1, Set, Rac1) in young and aged MuSCs measured by single-cell RNA seq (mean ± SEM). Two-tailed unpaired t-test analysis with Welch’s correction was used to determine the difference in gene expression between young and aged MuSCs. *, *** and **** represent statistical significance at p*≤*0.05, p*≤*0.001 and p*≤*0.0001 respectively.

**Fig. S4.**
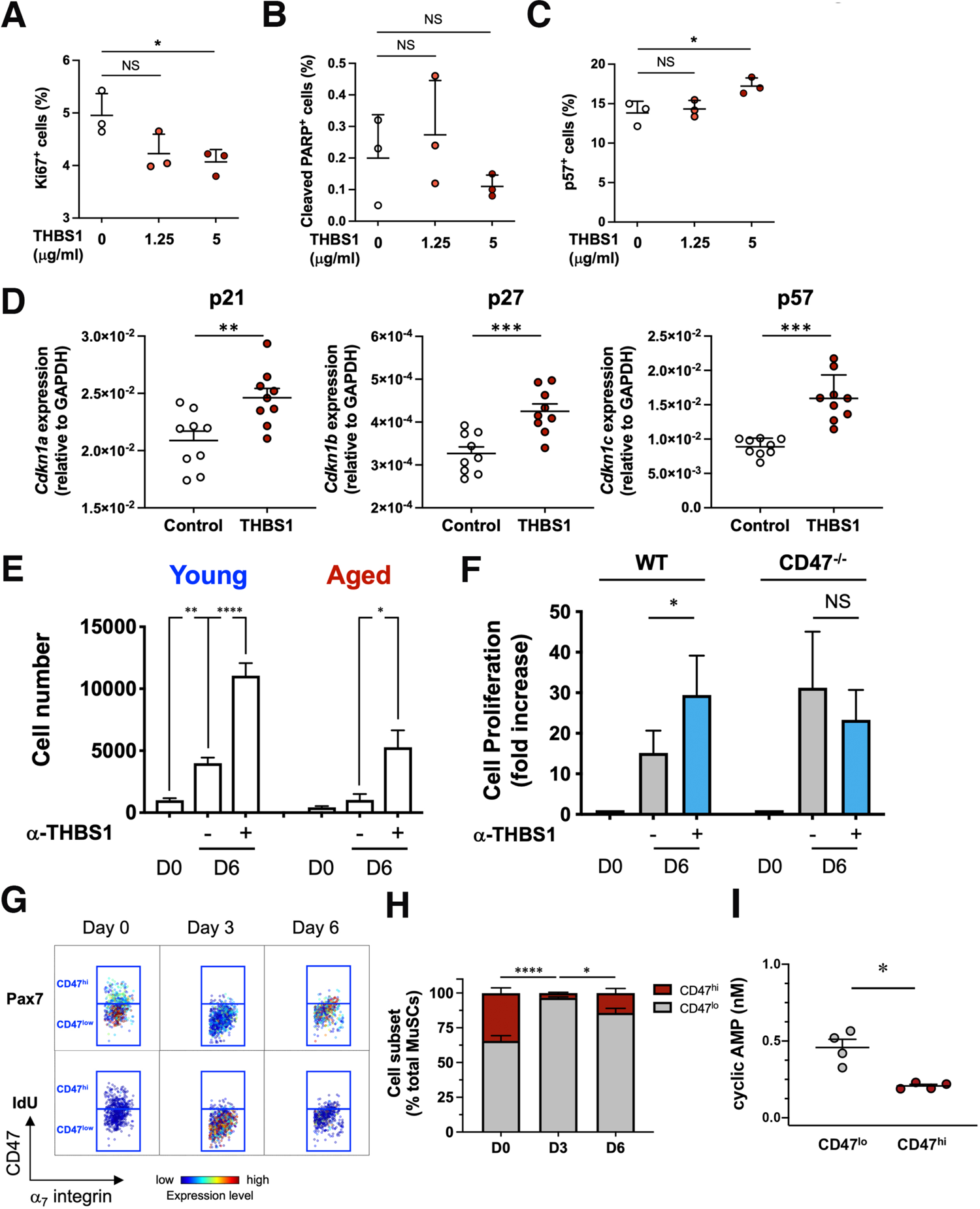
**(A-C)** The scatter plots show the fraction of ki-67^+^(A), cleaved PARP^+^(B), p57^+^(C) MuSCs, measured by CyTOF analysis in response to a 6-day treatment of young MuSCs with increasing doses of thrombospondin-1 (THBS1) (mean ± SD, n= 3 young mice). One-way ANOVA analysis with Tukey’s correction for multiple comparisons was used to determine difference in the fraction of ki-67^+^(A), cleaved PARP^+^(B), p57^+^(C) cells between untreated and THBS1 treated young MuSCs. **(D)** Expression levels of *Cdkn1a(p21)*, *Cdkn1b(p27)*, *Cdkn1c(p57),* measured by RT-qPCR in sorted MuSCs treated with THBS1 for 21h (mean ± SEM, n=9 replicates, 3 independent experiments). Two-tailed paired t test was performed to determine the difference in gene expression levels between THBS1 treated MuSCs and control treated MuSCs. **(E)** *α*_7_integrin^+^/CD34^+^ MuSCs sorted from young and aged mice were cultured in growth media for 6 days on biomimetic hydrogels in the presence (+) or absence (-) of a blocking antibody to THBS1 (*α*-THBS1) and proliferation was quantified by cell count (mean ± SEM from n=7 young replicates, n=3 aged replicates). Two-way ANOVA analysis with Tukey’s correction for multiple comparisons was used to determine difference between conditions in young and aged MuSCs. **(F)** MuSCs sorted from young wild type and CD47^-/-^ mice were cultured in growth media for 6 days on biomimetic hydrogels in the presence (+) or absence (-) of a blocking antibody to THBS1 and proliferation was quantified by cell count (mean ± SEM from n= 4 young replicates, n=4 aged replicates). Two-way ANOVA analysis with Tukey’s correction for multiple comparisons was used to determine difference between conditions in wild type and CD47^-/-^ MuSCs. **(G)** CyTOF analysis of young hindlimb muscle showing the dynamics of *α*_7_integrin^+^**/**CD34^+^ MuSC subsets during an injury time course. Representative biaxial dot plots of CD47 (y axis) by *α*_7_ integrin (x axis) colored by channel shows Pax7 expression (upper panels) and IdU incorporation (lower panels) within CD47^hi^ and CD47^lo^ MuSC subsets during the injury time course (Day 0, Day 3, Day 6). **(H)** Stacked column graph shows the proportion of CD47^lo^ (gray) and CD47^hi^ (dark red) MuSCs during the injury time course in (G) (mean ± SEM, n=9 mice, 3 independent experiments). Two-way ANOVA analysis with Tukey’s correction for multiple comparisons was used to determine difference in the proportion of CD47^lo^ and CD47^hi^ MuSCs between the different time points. **(I)** The scatter plot shows the cyclic AMP levels in sorted CD47^lo^ and CD47^hi^ MuSC subsets (mean ± SEM, n=4 mice, 2 independent experiments). Paired t test was performed to determine the difference in cAMP levels between the CD47^lo^ and CD47^hi^ MuSC subsets. *, **, *** and **** represent statistical significance at p*≤*0.05, p*≤*0.01, p*≤*0.001 and p*≤*0.0001 respectively.

**Fig. S5.**
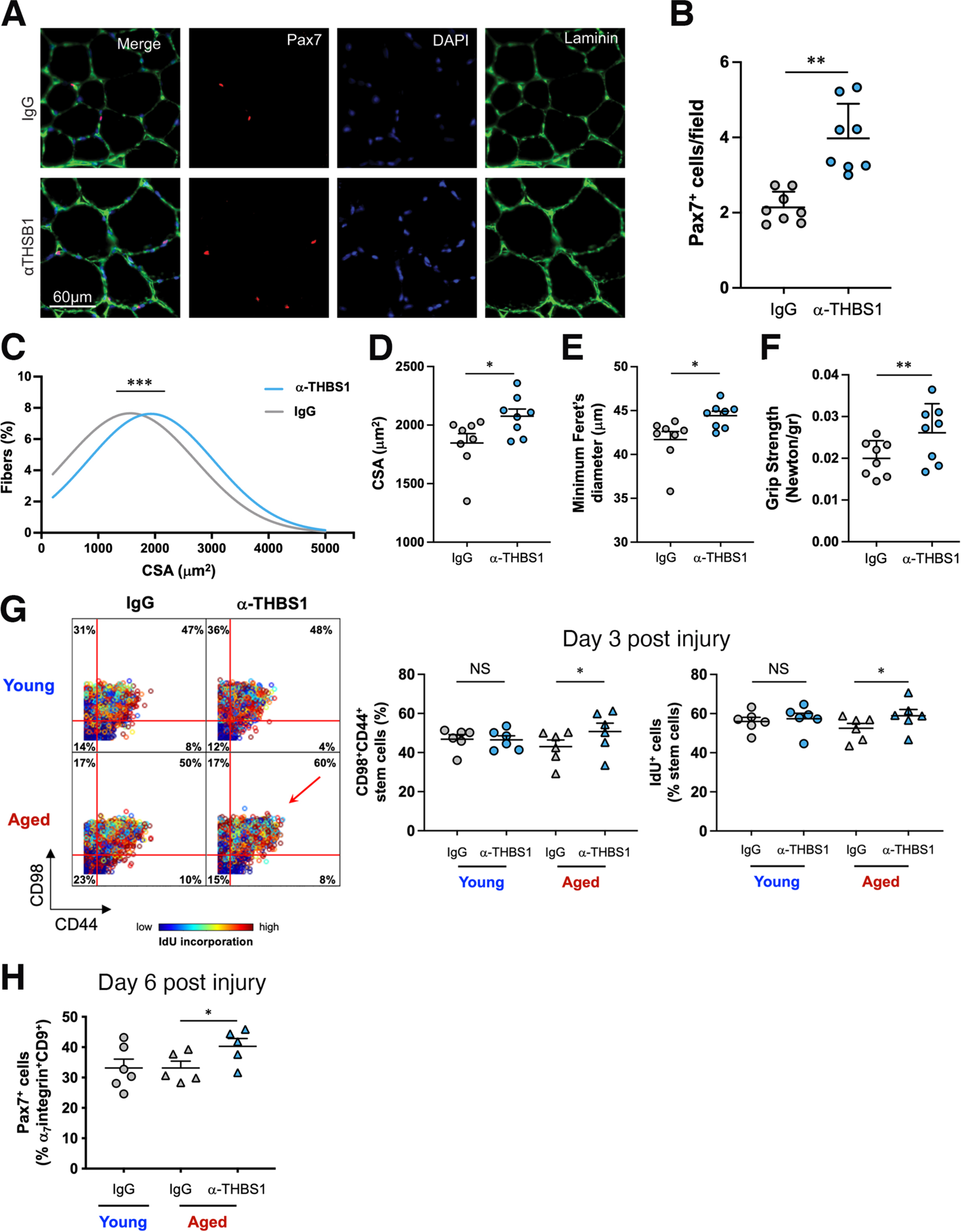
**(A-F)** Young Pax7^CreERT2^;Rosa26-LSL-Luc mice treated with tamoxifen (TAM) were pretreated (three times, at two-days intervals) by intramuscular injection in the TA muscle with a blocking antibody to THBS1 or IgG control (contralateral leg). Grip strength was measured and hindlimb muscle tissue was collected for histology 14 days after the end of treatment (n = 8 mice per condition, 2 independent experiments). **(A)** Immunohistochemistry analysis of TA muscles stained for PAX7 (red) and laminin (green) and counterstained with DAPI (blue). **(B)** Quantification of data in (A). The scatter plot shows the number of Pax7^+^ cells/field in IgG treated (gray) and *α*-THBS1 treated (light blue) TA muscles 14 days after the end of treatment (mean ± SEM). Two-tailed paired t-test was used to determine the difference in the number of Pax7^+^ cells between IgG control and *α*-THBS1 treated TA muscles. **(C)** Myofiber cross sectional area (CSA) was quantified in IgG treated (gray) and *α*-THBS1 treated (light blue) TA muscles and curve fitting was performed (mean ± SEM). Chi-square test was performed to determine the difference in the CSA distribution between IgG control and *α*-THBS1 treated TA muscles. **(D)** The scatter plot shows the mean CSA in sectioned IgG treated (gray) and *α*-THBS1 treated (light blue) TA muscles 14 days after the end of treatment (mean ± SEM). One-tailed paired t-test was used to determine the difference in mean CSA between IgG control and *α*-THBS1 treated young TA muscles. **(E)** The scatter plot shows the minimum fiber Feret’s diameter in IgG treated (gray) and *α*-THBS1 treated (light blue) TA muscles 14 days after the end of treatment (mean ± SEM). One-tailed paired t-test was used to determine the difference in the minimum fiber Feret’s diameter between IgG control and anti-THBS1 treated young TA muscles. **(F)** Scatter plot shows grip strength measurements, normalized to mouse weight, in *α*-THBS1 treated muscles and IgG treated contralateral muscles (mean ± SEM). Two-tailed paired t-test was used to determine the difference in grip strength between IgG control and *α*-THBS1 treated muscles. **(G, H)** Young and aged mice were treated (three times, at two-day intervals) with a blocking antibody to THBS1 or IgG control (contralateral leg) prior to notexin injury and hindlimb muscle tissue was collected for CyTOF analysis 3 days (G) or 6 days (H) post injury. **(G)** (Left) Representative biaxial dot plots of CD98 by CD44 colored by channel, show IdU incorporation in activated MuSCs, defined by co-expression of surface markers CD98 and CD44 (upper right quadrant). (Middle) The scatter plot shows the proportion of CD98^+^/CD44^+^ activated MuSCs (mean ± SEM, n=6, 2 independent experiments). One-tailed paired t-test was used to determine the difference in the abundance of CD98^+^/CD44^+^ MuSCs between IgG control and anti-THBS1 treatment in young or aged samples. (Right) The scatter plot shows the proportion of IdU^+^ cells in the MuSC population (mean ± SEM, n=6, 2 independent experiments). One-tailed paired t-test was used to determine the difference in the abundance of IdU^+^ MuSCs between IgG control and anti-THBS1 treatment in young or aged samples. **(H)** Scatter plot shows the fraction of Pax7^+^ cells in the Live/Lineage^-^/ *α*_7_integrin^+^/CD9^+^ cell population from young control mice and aged control and treated mice at day 6 post injury (mean ± SEM, n=6, 2 independent experiments). Paired t-test was used to determine the difference in the abundance of MuSCs between IgG control and *α*-THBS1 treatment in aged samples. *, ** and *** represent statistical significance at p*≤*0.05, p*≤*0.01 and p*≤*0.001 respectively.

